# PDIM drives the transition from repairable to catastrophic EsxA-mediated vacuole membrane damage during *Mycobacterium marinum* infection

**DOI:** 10.64898/2026.05.27.728140

**Authors:** Céline Michard, Angélique Perret, Mélanie Foulon, Hendrik Koliwer-Brandl, Hubert Hilbi, Thierry Soldati

## Abstract

During infection, pathogenic mycobacteria damage the membrane of the *Mycobacterium*-containing vacuole (MCV), triggering host repair responses that preserve an intracellular permissive niche before bacteria escape to the cytosol at later stages. How the MCV transitions from repairable injury to catastrophic rupture remains poorly understood. Here, we dissected the respective contributions of the ESX-1 effectors EsxA/EsxB and the cell-envelope lipid phthiocerol dimycocerosate (PDIM) during *Mycobacterium marinum* infection of the amoeba *Dictyostelium discoideum*. Combining host reporters for membrane damage and repair with single and double bacterial mutants, we show that EsxA/EsxB and PDIM are both required for full intracellular virulence but act at distinct stages of MCV damage progression. Loss of EsxA/EsxB strongly reduced recruitment of ESCRT and autophagy reporters to the MCV, demonstrating that EsxA initiates repairable membrane lesions. In contrast, PDIM-deficient bacteria retained the ability to recruit early damage reporters and ESCRT machinery but showed reduced autophagy-associated repair, failed to efficiently acquire cytosolic perilipin coating, and remained largely confined within the MCV. Genetic disruption of host autophagy restored cytosolic access and intracellular growth of PDIM-deficient bacteria, indicating that PDIM is specifically required to overcome host repair capacity rather than to initiate damage. Importantly, the sequential requirement for EsxA and PDIM was conserved during infection of murine microglial BV-2 cells. Remarkably, PDIM-defective mutants induced lysenin recruitment, a reporter of sphingomyelin exposure, but progression to autophagy engagement was strongly decreased. Together, our results support a two-step model in which EsxA initiates MCV membrane damage, while PDIM amplifies these lesions towards catastrophic rupture, enabling escape to the cytosol and dissemination.

## Introduction

*Mycobacterium tuberculosis* (Mtb) is the causative agent of tuberculosis, a severe pneumonia and the deadliest infection caused by a single bacterial pathogen worldwide, killing around 1 million persons each year (1). *Mycobacterium marinum* (Mm) is a close relative of Mtb (2, 3), inducing a tuberculosis-like disease in fish and amphibians. Mm is also an opportunistic pathogen for animals and humans, causing granulomatous skin lesions similar to the pulmonary lesions induced by Mtb (4). Mm and Mtb share most of their virulence factors (5, 6), among which the region of difference 1 (RD1) a locus carrying the genes encoding the ESX-1 type VII secretion system and its effectors. Deletion of RD1 is the key feature of the *M. bovis* BCG (Bacillus Calmette-Guérin) vaccine strain against Mtb (7).

The similarity between Mtb- and Mm-induced lesions results from their almost identical intracellular infection cycles (8, 9). Indeed, both Mtb and Mm are primarily phagocytosed by the macrophages of their hosts and subsequently reside in a vacuole called MCV for *Mycobacterium*-containing vacuole. The MCV transiently acquires early endosomal markers before further inhibition of the maturation and lysosomal degradation processes by the bacteria (10, 11). Mm and Mtb rapidly damage the membrane of their MCV, triggering host membrane repair mechanisms that act to reseal the compartment (12, 13). Cycles of membrane damage/repair occur, allowing bacteria to grow within the vacuole. At a later stage, the MCV fully ruptures and mycobacteria access the host cytosol where they continue to replicate before being released to the environment, enabling dissemination of the infection to naïve cells (14, 15). This infection cycle is also conserved in the environmental amoeba *Dictyostelium discoideum* (Dd), making this model organism a convenient system to study Mm molecular mechanisms.

Dd is a professional phagocyte that shares similar cell autonomous defence pathways with macrophages (8, 16). In its natural environment, Dd feeds on bacteria and therefore possesses a broad arsenal of anti-microbial mechanisms that parallel those of mammalian macrophages and neutrophiles (17, 18). In the last decades, Dd has also become more easily genetically tractable, making it a powerful model to study host-pathogen interactions, including those involving *Legionella pneumophila*, *Mycobacterium spp.*, *Vibrio cholera*, as well as yeast and fungi (9, 16).

During Mtb/Mm infection, cycles of membrane damage/repair represent a critical step in both macrophages and Dd. Due to their importance, several groups have investigated their mechanisms, highlighting a broadly conserved process. To induce MCV damage, pathogenic mycobacteria exploit changes in membrane composition that occur during the MCV maturation process. Indeed, sterol-rich microdomains are required for efficient membrane damage induction by Mm (19, 20). OSBP proteins, which exchange phosphatidylinositol-4-phosphate for sterol at MCV-endoplasmic reticulum contact sites, likely help sustain a high sterol concentration at the MCV (21). Membrane damage then leads to calcium leakage from the MCV contributing to the activation of the host repair response, including engagement of the endosomal sorting complex required for transport (ESCRT) and components of the autophagy machinery (22, 23). Both pathways are essential to maintain MCV integrity and for successful bacterial replication (11, 13, 22, 24). In the macrophage/Mtb model, cooperation between ESCRT and autophagy involves galectin-3 (25). In contrast, in the Dd/Mm model, the E3 ubiquitin ligase TrafE coordinates these two pathways (26), downstream of the calcium sensor PefA, an Alg2 ortholog (23). Recently, it has been proposed that during Mtb infection of macrophages, autophagy-mediated membrane repair can be either ESCRT-dependent or ESCRT-independent (22, 27). Moreover, in the macrophage/Mtb model, an additional repair pathway involving membrane wound “plugging” by stress granules has been identified (28, 29).

In recent years, it has been recognized that membrane atg8ylation is central not only to canonical autophagy, but also to a broader range of homeostatic processes that encompass membrane repair and remodelling pathways in cells subjected to microbe-induced or sterile stress and membrane damage (22, 30-32). In Dd, ATG8ylation also occurs but it has not yet been associated with cellular processes other than canonical autophagy. In this paper, we use the terms “autophagy” or “autophagy-related” to describe the repair mechanisms involving the recruitment of at least some components of the autophagy machinery (11, 13).

Membrane damage/repair cycles require a fine balance between bacterial damage induction and host repair activity. Mtb or Mm carrying an RD1 deletion show a drastically reduced ability to induce MCV membrane damage (33) and remain confined within their vacuole, with minimal replication (34). In contrast, knockout of the ESCRT repair machinery in Dd leads to premature escape of Mm from its MCV to the cytosol, triggering its degradation via xenophagy (13, 23). Incapacitation of the autophagy repair machinery in macrophages or Dd also results in early vacuole escape, but in this case favours Mtb/Mm replication because xenophagy is also impaired (13, 35). Membrane damage at the MCV therefore appears to be tightly controlled. However, at later stages of infection, Mtb and Mm ultimately access the host cytosol (14, 36), which is a prerequisite for further growth and dissemination. A key unresolved question is how pathogenic mycobacteria transition from maintaining MCV integrity to fully escaping to the cytosol. One hypothesis is that bacteria sequentially induce two categories of membrane damage: initial lesions that can be efficiently repaired by host machineries, followed by catastrophic damage that exceeds repair capacity and leads to cytosol escape. Alternatively, a single type of membrane damage may progressively enlarge over time until repair mechanisms become insufficient. To understand this crucial compartment transition, one needs to consider the various effectors implicated in how pathogenic mycobacteria rupture the MCV membrane.

Twenty years ago, EsxA (also known as ESAT-6 for 6 kDa early secretory antigenic target) was identified as a major Mtb/Mm virulence factor. Its absence in *M. bovis* BCG and Mtb/Mm ΔRD1 strains is responsible for their attenuation (7, 37). EsxA forms a dimer with its chaperone EsxB (also called CFP-10 for 10 kDa culture filtrate protein) (38). This complex is secreted through the ESX-1 system and participates in the secretion of other ESX-1 substrates (39-43). EsxA has been shown to modulate several host cellular pathways (44, 45), but its predominant function is to induce membrane damage during infection (12, 33, 46). Even though Mtb/Mm arrest phagosome maturation, a slight and transient acidification occurs (47). Secretion of EsxA/EsxB into this mildly acidic environment leads to dissociation of the dimer (46). It has been proposed that EsxA, like other (chole)sterol-dependent toxins, inserts into sterol-rich microdomains of the host membrane, generating pores that contribute to the formation of a permissive replication niche (12, 20, 48, 49). This model was challenged by Conrad *et al.* who identified a requirement for direct bacteria/ host membrane contact to induce membrane damage (50). Moreover, Mm transposon mutants defective in EsxA secretion were still able to induce some level of phagosome damage, suggesting that EsxA alone is insufficient to explain ESX-1-mediated MCV disruption (51). Alternative modes of EsxA membrane interaction have also recently been reported (52) and the precise molecular mechanism is still to be elucidated.

Membrane damaging activity has also been reported for phthiocerol dimycocerosates (PDIM) (53-56). PDIM is a noncovalently bound lipid found in the cell envelope of pathogenic mycobacteria (57). It can be shed from the bacterial surface and spread within and between host cells during infection (58). The mechanism by which PDIM contributes to membrane damage remains unclear but based on *in vitro* liposome assays, Augenstreich *et al.* proposed that PDIM enhances EsxA activity (56, 59). Altogether, intracellular pathogenic mycobacteria appear to possess multiple mechanisms to induce membrane damage, yet the overarching molecular basis behind and particularly for cytosol escape remains unresolved.

Here, by building on current knowledge, we sought to dissect in detail how damage/repair cycles operate during Mm infection. We generated single and double Mm mutants, carrying deletions in RD1, *esxA*/*esxB* and/or *tesA*, the latter abolishing PDIM production. By monitoring intracellular growth of these Mm mutants in both Dd and mammalian BV-2 microglial cells, we found that loss of EsxA or PDIM does not result in equivalent attenuation of virulence. Analysis of repair machineries recruitment to their respective MCV, as well as monitoring of bacterial access to the cytosol, identified EsxA as an inducer of repairable membrane damage. In contrast, PDIM was required to amplify this damage and enable progression towards catastrophic damage and ultimately Mm cytosol escape and dissemination. Our results demonstrate that PDIM is absolutely required for the completion of the process but can act only once the initial EsxA-mediated damage have formed. This two-step mechanism was also observed in mammalian cells, indicating a striking evolutionary conservation of bacterial virulence and host defence pathways.

## Results

### PDIM and EsxA are differently required to achieve full Mm virulence in Dd

To decipher in more detail the mechanisms of membrane damage/repair cycles occurring during pathogenic mycobacteria infection, we used a set of Mm mutants deleted for bacterial components known to induce membrane damage (Fig 1a) (33, 50, 56). Mm ΔRD1 and Mm ΔCE (or Mm Δ*esxA/esxB*) mutants allowed us to differentiate the contributions of the ESX-1 type VII secretion system and the EsxA/EsxB dimer, respectively. To study the role of PDIM, we generated a Mm mutant deleted for the type II thioesterase *tesA* gene (60, 61), which abolishes synthesis of PDIM and PGL. This experimental strategy is congruent with the previous studies on the contributions of PDIM to Mtb/Mm-induced damage that were performed using strains carrying mutations affecting both PDIM and PGL syntheses. The Mm *tesA* deletion was also introduced into the Mm ΔRD1 and Mm ΔCE strains to generate double Mm mutants lacking either the ESX-1 system and PDIM/PGL synthesis, or the EsxA/EsxB dimer and PDIM/PGL synthesis, respectively. Secretion assays were performed to confirm that Mm wild-type (WT) and Mm Δ*tesA* were able to produce and secrete the EsxA/B dimer, in contrast to the mutants with the ΔRD1 or ΔCE deletions (Fig S1a). The disruption of PDIM/PGL synthesis in all Mm mutants carrying a *tesA* deletion was also confirmed by thin layer chromatography of total lipid extracts (Fig S1b, S1c). Finally, the *in vitro* growth of the Mm mutants was examined, and their doubling time were similar (Fig S1d), validating that there is no large fitness cost in broth due to the mutations.

**Fig 1.**
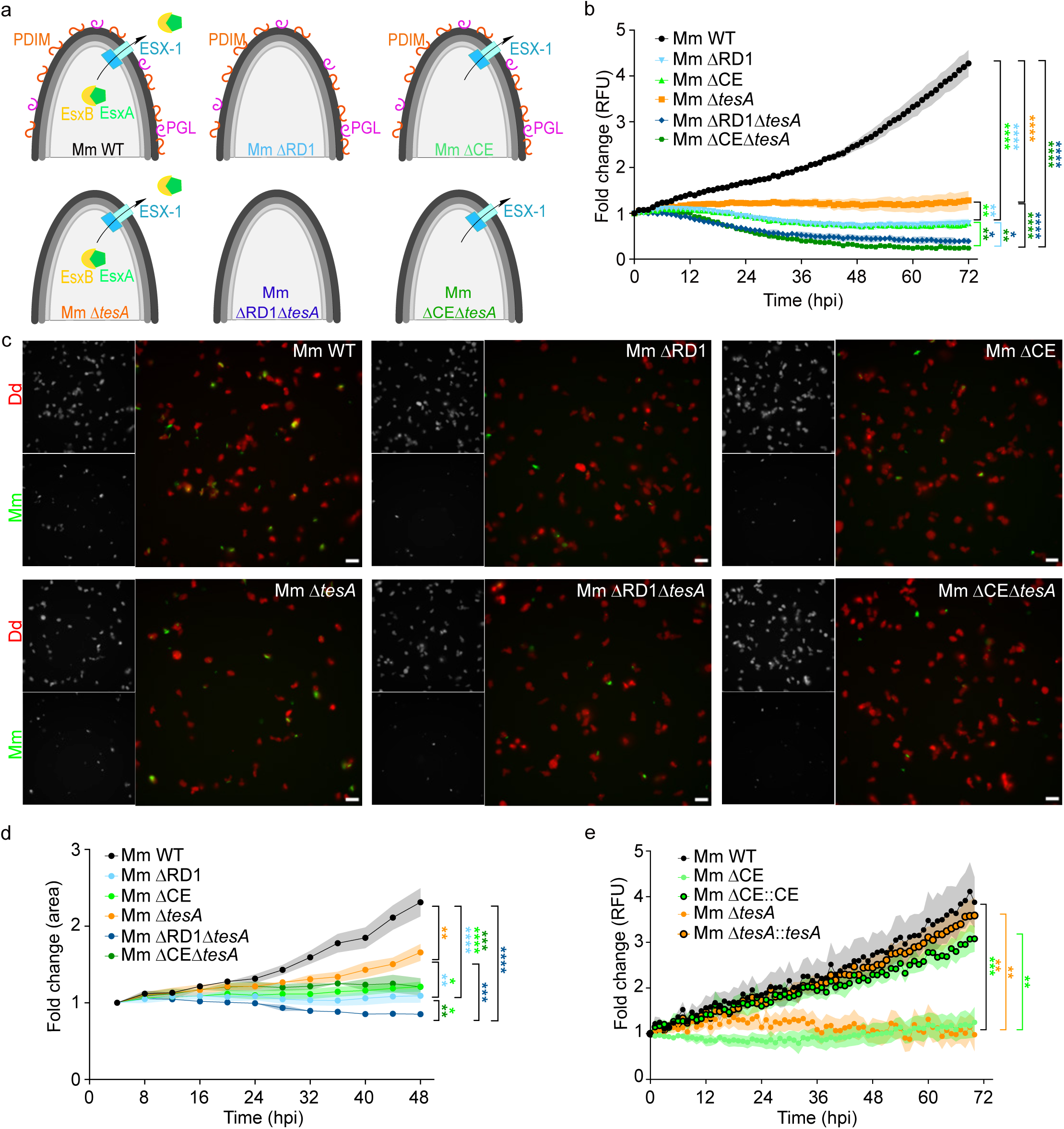
PDIM and EsxA/EsxB Mm mutants are differently attenuated in Dd. (a) Scheme presenting the main characteristics of the Mm mutants used in this study. (b) Dd were infected with GFP-expressing Mm mutants and green fluorescence intensity was monitored for 72 hours by plate reader. Data are represented as the average fold change ± SEM (n = 3, N = 3, two-way ANOVA with Fisher’s LSD test). RFU, relative fluorescence unit. (c-d) mCherry-expressing Dd infected with GFP-expressing Mm mutants and monitored by high-throughput, high-content microscopy for 48 hours. (c) Representative pictures of infected cells at 48 hpi. Scale bar, 20 µm. (d) Growth curves of the Mm mutants measured as area sum of the GFP signal every 4 hours and represented as the mean of fold change ± SEM (9 pictures per well, n = 3, N = 3, two-way ANOVA with Fisher’s LSD test). (e) Dd were infected with GFP-expressing Mm mutants and complemented strains. Green fluorescence intensity was measured every hour for 72 hours by plate reader. The graph represents the average fold change ± SEM (n = 3, N = 4, two-way ANOVA with Fisher’s LSD test). RFU, relative fluorescence unit.

The intracellular growth of all the strains in Dd was then assessed to evaluate the respective contributions of ESX-1, EsxA/EsxB and PDIM/PGL to Mm virulence (Fig 1b-d). The intracellular growth of Mm GFP was monitored for 72 hours with a plate reader, measuring fluorescence intensity as a proxy for bacteria load (Fig 1b). The intracellular growth of each Mm mutant was significantly reduced compared to Mm WT, while their *in vitro* growth was similar (Fig S1d), demonstrating that the observed attenuations were specific to infection. Deletion of the genes encoding the ESX-1 system (Mm ΔRD1) or the EsxA/EsxB dimer (Mm ΔCE) resulted in identical levels of attenuation, confirming that the EsxA/EsxB dimer is the main virulence factor secreted by the ESX-1 system. Mm Δ*tesA* displayed an intermediate intracellular growth compared to Mm WT and Mm ΔCE, indicating that the absence of PDIM/PGL attenuates Mm virulence, although to a lesser extent than the absence of the EsxA/EsxB dimer. Finally, the two double Mm mutants (Mm ΔRD1Δ*tesA* and Mm ΔCEΔ*tesA*) showed similar and strongest attenuation among all mutants, indicating an additive effect of the two deletions.

The intracellular attenuation was not due to a defect in uptake, as a phagocytosis assay showed similar uptake for all Mm strains, with even a slightly higher uptake for Mm ΔRD1 compared to Mm WT (Fig S1e). In addition, the plate reader experiments confirmed the absence of an uptake defect for each Mm mutant, with initial fluorescence values ranging from 1.1- to 1.7-fold higher than Mm WT on average (Fig S1f).

To further confirm the intracellular attenuation of the Mm mutants, high-throughput microscopy experiments were performed (Fig1c,d). Infected Dd were seeded, and multiple images of each infection were acquired every 4 hours for 48 hours using an automated microscope. At the end-point of 48 hours post-infection (hpi), the bacteria load differed depending on the Mm strain used, with Mm WT showing higher bacteria levels compared to all Mm mutants, and Mm Δ*tesA* presenting an intermediate phenotype (Fig 1c). High-content quantification of the bacteria load over time (Fig 1d) confirmed that all the Mm mutants displayed an attenuation compared to Mm WT, with a hierarchy similar to that obtained with the fluorescence plate reader assay, except for the double Mm mutant ΔCEΔ*tesA* which showed a level comparable to the single Mm ΔCE and Mm ΔRD1. Importantly, Mm Δ*tesA* again showed an intermediate attenuation, while the growth of Mm ΔCE was more strongly impaired, demonstrating that PDIM/PGL and EsxA/EsxB are both required but contribute differently.

To validate the specificity of the phenotypes observed for the single deletion Δ*esxA/esxB* and Δ*tesA*, complemented Mm strains were generated using integrative plasmids. PDIM and PGL synthesis were restored in the Mm Δ*tesA*::*tesA* strain (Fig S1b, S1c). However, we did not detect WT levels of the EsxA/EsxB dimer in the Mm ΔCE::CE strain, despite using a complementation strategy similar to that originally described for this mutant (Fig S1a) (37). Prolonged exposure of the western-blot membrane showed a small amount of EsxB in the bacterial lysate of Mm ΔCE::CE. Because EsxA and EsxB are expressed as a dimer, we assume that low levels of Mm EsxA are also produced but not detected by the anti-Mtb EsxA antibody. We then proceeded to test the virulence of both complemented strains. First, the integration of the plasmids did not affect the *in vitro* growth of the Mm strains (Fig S1d). But convincingly, both complementation plasmids significantly increased the intracellular growth of their corresponding Mm mutants to levels comparable to Mm WT (Fig 1e), confirming that the observed intracellular attenuation resulted from the specific deletions.

Moreover, to demonstrate that the attenuation of Mm Δ*tesA* was specifically linked to the absence of PDIM rather than PGL, we generated a Mm Δ*pks15/1* mutant lacking a polyketide synthase specifically involved in PGL synthesis. Thin layer chromatography of total lipid extracts confirmed the absence of PGL in this Mm mutant without affecting PDIM levels (Fig S2a,b). The Mm Δ*pks15/1* mutant did not show any intracellular attenuation in the plate reader assay, in contrast to Mm Δ*tesA* (Fig S2c), indicating that the phenotype observed for Mm Δ*tesA* results solely from the absence of PDIM.

Altogether, these results indicate that both EsxA/EsxB and PDIM are required to achieve full virulence of Mm in Dd but contribute to different extents and likely in different manner.

### The ESCRT and autophagy machineries are extremely dynamic and are differentially recruited during MCV damage

Both ESCRT and autophagy machineries are recruited to damaged MCVs, however they have never been visualized simultaneously in live cells (11, 13, 22, 24). We developed a Dd cell line stably expressing GFP-Vps32, a subunit of ESCRT-III, and mCherry-Atg8A, a homologue of the mammalian LC3, from chromosomal loci, allowing simultaneous live imaging of both ESCRT and autophagy pathways (62). After infection with Mm WT, the cells were confined under an agarose sheet to reduce their motility and monitored for several hours by spinning-disk confocal microscopy. Three types of reporter associations were observed (Fig 2a-c and S3a). Few MCVs recruited only GFP-Vps32 (Fig 2a), while the majority of MCVs were associated with mCherry-Atg8A at least at one time point (Fig 2b). Both reporters were present as dots or patches, and in some cases appeared to cover the entire MCV. Both reporters were most often recruited to the same MCV (Fig 2c), but mainly in juxtaposed, non-overlapping positions. These results are consistent with previous observations (11, 13).

**Fig 2.**
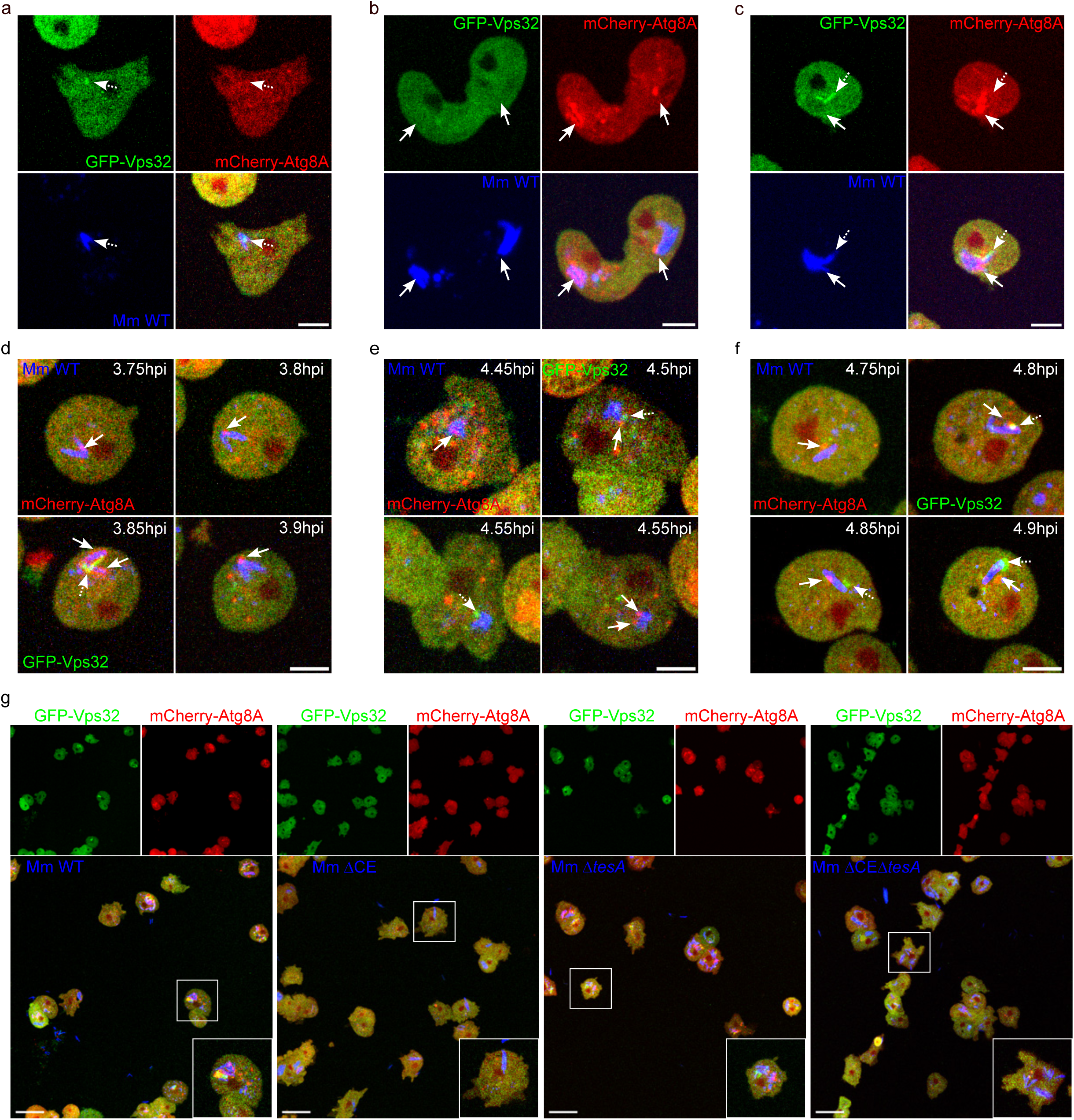
ESCRT and autophagy are both dynamically recruited at the MCVs. (a-f) Dd expressing both GFP-Vps32 and mCherry-Atg8A was infected with EBFP-expressing Mm WT and followed by live imaging for several hpi with pictures taken every 5 minutes. Representative maximum projections from 2 sections show specific GFP-Vps32 (a) (dotted arrow), mCherry-Atg8A (b) (full arrow), or double GFP-Vps32 (dotted arrow)/ mCherry-Atg8A (full arrow) (c) recruitment at the MCV. Scale bar, 10 µm. (d-f) Examples of the dynamic of ESCRT and autophagy recruitment at three independent MCVs are shown using maximum projections from 2 sections. A dotted arrow identifies a GFP-Vps32 recruitment at the bacteria and a full arrow corresponds to the presence of mCherry-Atg8A at the MCV. Scale bar, 10 µm. (g) Infections of Dd expressing both GFP-Vps32 and mCherry-Atg8A by EBFP-expressing Mm mutants were monitored by live imaging for several hpi. Images are representative maximum projections at 4 hpi. Scale bar, 20 µm.

Both machineries were extremely dynamic and, with one exception, all MCVs tracked for at least 10 minutes recruited one of the reporters at some point (Fig 2d-f and S3a). Figure 2d-f illustrate this dynamic behaviour around three representative MCVs. In Figure 2d, a single mCherry-Atg8A dot was initially observed and slightly changed localization over time, before the MCV became fully covered by mCherry-Atg8A together with an extended GFP-Vps32 patch. Shortly after, this signal returned to a single mCherry-Atg8A dot. In Figure 2e, a mCherry-Atg8A dot changed position within minutes while a GFP-Vps32 dot appeared on the opposite side. The autophagy reporter then disappeared from the MCV as the ESCRT component persisted and relocalized, before being replaced again by two mCherry-Atg8A dots. In Figure 2f, both localization and structure size vary markedly, with transitions from small patches to larger structures for both reporters, and changes in localization around the MCV. Overall, the duration of each reporter recruitment, as well as their partner (dots, patches, coating), varied between MCVs (Fig S3). This result highlights the dynamic and heterogeneous recruitment of ESCRT and autophagy machineries at the MCV.

We then monitored in live microscopy the recruitment of GFP-Vps32 and mCherry-Atg8A at the MCVs containing Mm ΔCE, Mm Δ*tesA* and Mm ΔCEΔ*tesA* (Fig 2g). Each Mm mutant displayed a distinct pattern of reporter association compared to Mm WT. Mm ΔCE and Mm ΔCEΔ*tesA* showed almost no recruitment of ESCRT and autophagy reporters to their MCVs. In contrast, cells infected with Mm Δ*tesA* displayed patches of GFP-Vps32 and mCherry-Atg8A around the MCVs, although to a lesser extent than Mm WT. Therefore, both ESCRT and autophagy machineries are dynamic repair pathways that are recruited specifically, but heterogeneously, to damaged MCVs.

### Repairable MCV damage depends mainly on EsxA while escape to the cytosol requires both PDIM and EsxA

As Mm ΔCE, Mm Δ*tesA* and Mm ΔCEΔ*tesA* presented different levels of reporter recruitment in the double Dd cell line expressing both GFP-Vps32 and mCherry-Atg8A, we performed high-throughput, high-content time lapse microscopy over 36 hours to precisely quantify the individual recruitment of these two reporters for each Mm mutant (Fig 3a, 3b). During the first 12 hours of infection, 34-37% of the Mm WT MCVs were positive for GFP-Vps32, before decreasing to 18% at 36 hpi (Fig 3a). Mm Δ*tesA* showed the same GFP-Vps32 kinetic as Mm WT, with similar percentages of GFP-Vps32 positive MCVs at the different time points. In contrast, Mm ΔCE displayed significantly less ESCRT machinery recruitment at its MCVs, with only 4-7% of them positive for GFP-Vps32 over the 36 hpi, confirming the previous observations with the double reporter cell line (Fig 2e). The double Mm mutant ΔCEΔ*tesA* showed similar percentage of GFP-Vps32 recruitment as Mm ΔCE. Thus, the EsxA/EsxB dimer is required to induce ESCRT-repairable MCV membrane damage, and PDIM is dispensable.

**Fig 3.**
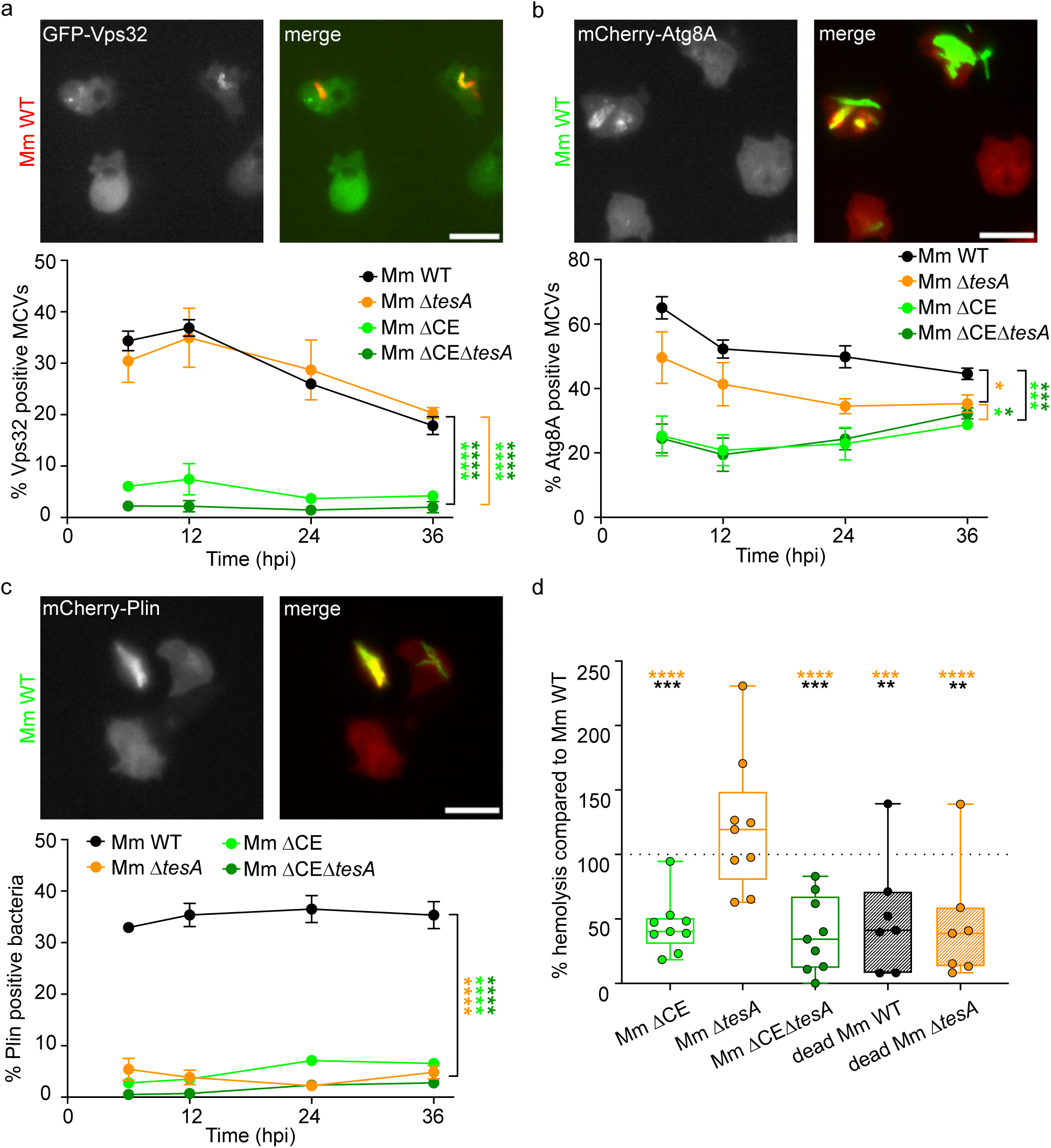
EsxA is necessary to recruit ESCRT and autophagy while PDIM allows cytosol access. (a-c) Dd expressing GFP-Vps32 (a), or mCherry-Atg8A (b), or mCherry-Plin (c) were infected with GFP- or mCherry- expressing Mm mutants and imaged live by high-throughput, high-content microscopy at the indicated time points. The images are representative pictures of the different marker recruitments at the MCV of Mm WT. Scale bar, 5 µm. The graphs represent the average ± SEM of the automated quantification of marker recruitment at the bacteria (9 pictures per well, n = 3-4, N = 3, two-way ANOVA with Fisher’s LSD test). (d) Haemolytic activity of Mm mutants and amikacin-killed Mm WT and Mm Δ*tesA* compared to Mm WT. The dotted line marks 100% and corresponds to Mm WT haemolysis. Each data point corresponds to one biological replicate (n = 1, N = 9, one-way ANOVA with Fisher’s LSD test).

The kinetic and extent of autophagy machinery recruitment appeared different than the ones for ESCRT (Fig 3b). Approximately 65% of Mm WT MCVs were positive for mCherry-Atg8A at 6 hpi, before a significant decrease to 52% at 12 hpi, and then a gradual decline to 45%. Mm ΔCE had a drastically reduced autophagy recruitment to its MCV, with 21-29% of MCVs positive for mCherry-Atg8A at all time points. Mm Δ*tesA* displayed an intermediate phenotype with 50% of its MCVs recruiting the autophagy machinery at 6 hpi, which then decreased to 35% at 36 hpi. As observed for ESCRT recruitment, the double Mm mutant ΔCEΔ*tesA* showed a phenotype similar to Mm ΔCE, suggesting that its attenuation is mainly due to the absence of the EsxA/EsxB dimer. Altogether, these results show that the EsxA/EsxB dimer triggers both ESCRT and autophagy recruitment to damaged MCVs. In contrast, absence of PDIM only partially reduces autophagy recruitment, likely because membrane damage is decreased in absence of PDIM. In other words, these data support a model in which autophagy repairs damage not resolved by ESCRT and PDIM plays a role in damage progression.

As previous studies showed that Mtb/Mm mutants lacking PDIM synthesis have reduced membrane damaging abilities (53-56) and because we also detected a decrease in autophagy recruitment around Mm Δ*tesA* MCVs, we hypothesized that PDIM participates in the progression from repairable to catastrophic damage, allowing bacteria escape to the host cytosol. To evaluate the ability of Mm mutants to access the cytosol, we used perilipin (Plin) as a diagnostic reporter to distinguish cytosolic from vacuolar bacteria. Plin is a lipid droplet stabilizer that also binds the hydrophobic mycobacterial cell wall once bacteria reach the cytosol in Dd and BV-2 microglial cells (63). We thus quantified the percentage of bacteria coated with Plin for each mutant as a read-out for cytosol escape (Fig 3c). As expected, about 30% of Mm WT accessed the host cytosol over the course of infection, with a slight increase over time. The cytosolic release of Mm ΔCE and Mm ΔCEΔ*tesA* was significantly reduced compared to Mm WT, with only 2-7% of bacteria becoming Plin-positive, indicating that the induction of repairable membrane damage by EsxA is required for efficient escape. Remarkably, the absence of PDIM also resulted in a strongly decreased percentage of bacteria being coated by Plin, with Mm Δ*tesA* reaching levels similar to the Mm ΔCE mutant, indicating that PDIM is necessary for damage progression and consequently cytosol escape.

Because the cell envelope of Mm Δ*tesA* is likely altered in absence of PDIM and PGL (60), we wondered whether the absence of Plin-binding to Mm Δ*tesA* was due to bacteria confinement within the MCV or to intrinsically low Plin-binding. To address this question, we developed an *in vitro* Plin-binding assay (Fig S4). Broth-grown Mm WT and Mm Δ*tesA* were incubated with cytosolic extracts from Dd expressing mCherry as control or mCherry-Plin, and protein binding to the bacteria was evaluated by microscopy (Fig S4a) and flow cytometry (Fig S4b,c). The fluorescent mCherry protein did not bind Mm WT or Mm Δ*tesA*. In contrast, mCherry-Plin decorated the Mm WT surface and induced a shift in fluorescence intensity during flow cytometry analysis. mCherry-Plin also bound the Mm Δ*tesA* mutant, with an apparently higher affinity compared to Mm WT. Therefore, the strong reduction in Plin-coating of Mm Δ*tesA* during Dd infection indicates that the absence of PDIM results in bacterial confinement within the MCV, and not to lower affinity for Plin. The fact that the double Mm ΔCEΔ*tesA* mutant had identical Plin-coating compared to Mm ΔCE and Mm Δ*tesA* but phenocopied only Mm ΔCE for ESCRT and autophagy recruitments, supports a hierarchical process in which EsxA initiates damage and PDIM is required for damage progression and cytosol escape.

Next, we decided to compare the results from the damage reporter recruitment to the haemolysis assay, a classic *in vitro* assay used to evaluate Mtb/Mm membrane damage ability (Fig 3d). Mm ΔCE and Mm ΔCEΔ*tesA* showed more than two-fold reduced haemolytic ability compared to Mm WT, whereas Mm Δ*tesA* was similar to the Mm WT control. As haemolytic activity appeared to correlate with EsxA secretion, we next tested dead Mm WT and Mm Δ*tesA*, killed by overnight incubation with amikacin to preserve their intact cell wall structure while arresting their metabolism. Dead Mm WT and dead Mm Δ*tesA* are unable to secrete proteins and showed significantly reduced haemolytic activity compared to Mm WT, confirming that haemolysis results from secreted proteins and mainly from EsxA.

Altogether, these results demonstrate that EsxA is the main mycobacterial factor inducing initial membrane damage at the MCV, consisting of repairable lesions that recruit both ESCRT and autophagy machineries. PDIM is also required for membrane damage and is crucial to enable progression towards catastrophic lesions and consequently, escape to the host cytosol. While EsxA damage activity is detected by the classical haemolysis assay, this *in vitro* test does not reflect the PDIM damaging capacity.

### ESCRT and autophagy impairment, but not altered MCV membrane composition, influence the intracellular growth of the Mm mutants

To better understand how EsxA and PDIM activities might be influenced by membrane composition and host repair pathways, we compared the intracellular growth of the Mm mutants under different conditions and in different Dd genetic backgrounds. It was previously shown that the MCV membrane composition, and specifically the presence of sterol-rich microdomains, is critical for EsxA to induce membrane damage (20). Deletion of the three Dd vacuolins (Dd Δ*vacABC*), homologues of the mammalian microdomain organizer flotillins, or the addition of methyl-β-cyclodextrin to Dd WT cells (Dd + CD), which extracts sterol from sterol-rich membranes, significantly decreased the intracellular load of Mm WT (Fig 4a, 4b, 4c). Importantly, the reduction in MCV damage resulted in confinement of Mm WT and thus phenocopied infection with the Mm ΔRD1 mutant. Under conditions that disrupt sterol-rich microdomains, the growth of Mm ΔCE (Fig 4a), Mm Δ*tesA* (Fig 4b) and Mm ΔCEΔ*tesA* (Fig 4c) was not further attenuated. Because Mm Δ*tesA* induces EsxA-mediated damage, a stronger growth restriction would have been expected upon disruption of sterol-rich microdomains. However, this effect may have been masked by the already substantial attenuation observed under these specific conditions. Therefore, these results indicate that disruption of sterol-rich microdomains is not detrimental for Mm mutants lacking EsxA/EsxB or probably PDIM, contrary to Mm WT.

**Fig 4.**
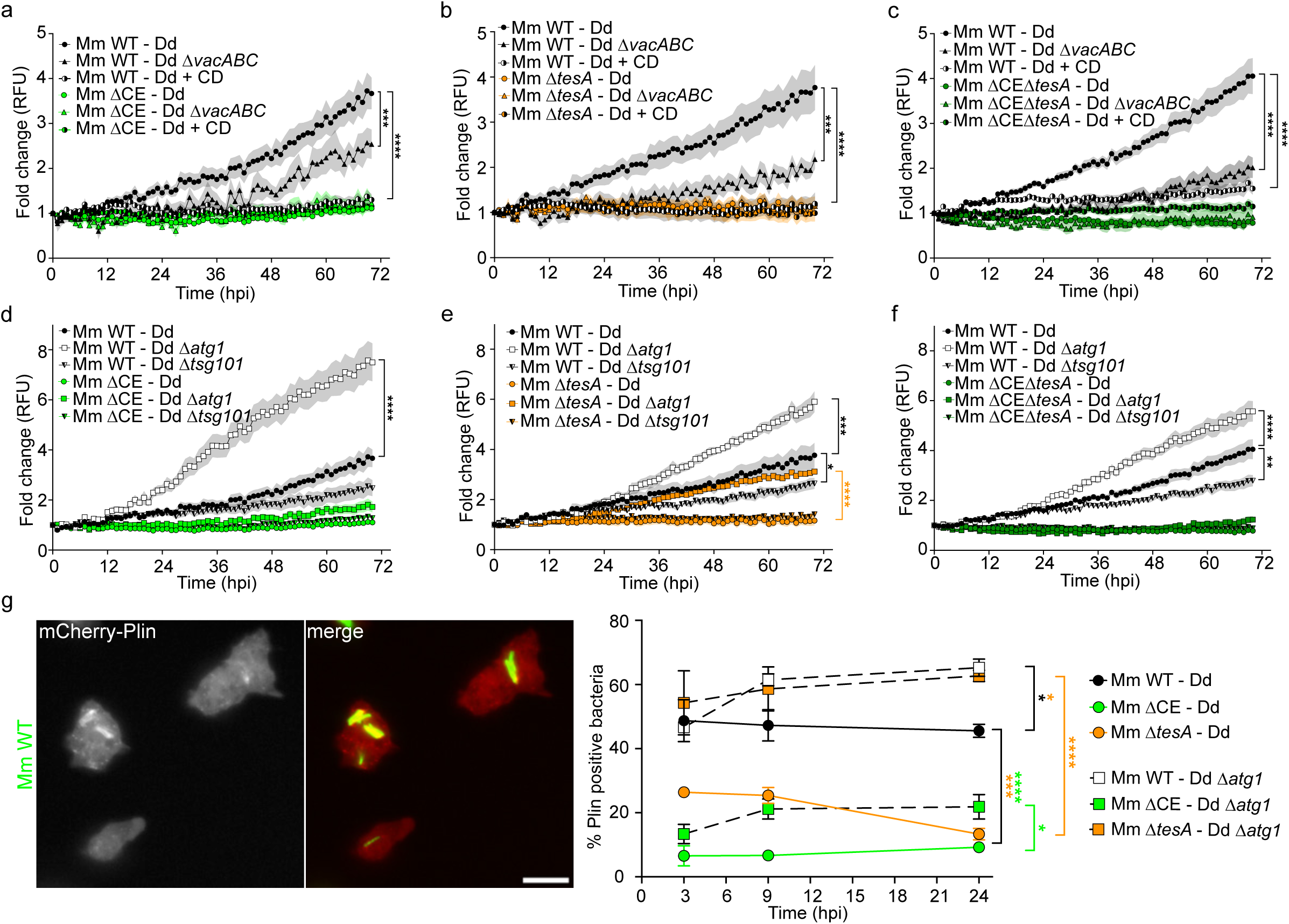
The absence of autophagy rescues the growth defect of the Mm Δ*tesA* mutant. (a-f) Different Dd backgrounds were infected with GFP-expressing Mm mutants. Green fluorescence was monitored for 72 hours by plate reader. (a), (b) and (c) compare the bacteria intracellular growth of Mm WT and Mm ΔCE, Mm Δ*tesA* or Mm ΔCEΔ*tesA*, respectively, into Dd WT (Dd), Dd deleted for vacuolins (Dd Δ*vacABC*) or in Dd in presence of 2 mM methyl-β-cyclodextrin (Dd + CD). (d), (e) and (f) present the intracellular growth of Mm WT and Mm ΔCE, Mm Δ*tesA* or Mm ΔCEΔ*tesA*, respectively, into Dd WT (Dd), Dd deleted for the ESCRT machinery (Dd Δ*tsg101*) or in Dd deleted for autophagy (Dd Δ*atg1*). Each graph represents the mean fold change ± SEM (n = 3, N = 4-5, two-way ANOVA with Fisher’s LSD test). RFU, relative fluorescence unit. (g) Dd and Dd Δ*atg1* expressing mCherry-Plin were infected with GFP-expressing Mm mutants and imaged live by high-content microscopy at several time points. The images are representative pictures of mCherry-Plin recruitment at the MCV of Mm WT. Scale bar, 5 µm. The graphs show the average recruitment ± SEM of the automated quantification of mCherry-Plin at the bacteria (9 pictures per well, n = 3, N = 3, two-way ANOVA with Fisher’s LSD test).

Next, we investigated the importance of functional ESCRT and autophagy pathways for the growth of Mm mutants. The autophagy machinery plays a dual role during Mm infection, contributing both to MCV membrane repair and to xenophagy-mediated restriction of cytosolic mycobacteria (11, 13, 26), two processes downstream of membrane damage. Deletion of the master regulator *atg1* blocks both autophagy-related pathways, and consequently, leads to premature cytosol escape of Mm WT and to its unrestricted growth (11). In contrast, defects in the ESCRT repair machinery (e.g. in Dd Δ*tsg101* or Dd Δ*alix*) result in attenuation of Mm WT, because premature escape to the cytosol leads to severe restriction by functional xenophagy (13, 23). A Mm strain drastically impaired in its damaging ability, such as Mm ΔRD1, does not escape prematurely from its vacuole in either host genetic background, and therefore, its intracellular load is not increased in Dd Δ*atg1* nor decreased in Dd Δ*tsg101* (11, 13). These two genetic backgrounds can thus be used as sensitive “tools” to evaluate bacteria membrane damaging capacity *in vivo*. As expected, in these conditions, Mm WT was attenuated in Dd Δ*tsg101* and grew unrestricted in Dd Δ*atg1* (Fig 4d, 4e, 4f), while absence of ESCRT or autophagy machineries did not impact the growth of Mm ΔCE (Fig 4d). Interestingly, Mm Δ*tesA* was differentially affected by the absence of each membrane repair machinery (Fig 4e). Mm Δ*tesA* was not further attenuated in Dd Δ*tsg101* compared to its growth in Dd, corroborating previous evidence that it does not reach the cytosol (Fig 3c) and is not further restricted by xenophagy. In contrast, disruption of autophagy pathways in Dd Δ*atg1* resulted in a marked increase of Mm Δ*tesA* growth, restoring bacteria loads to levels comparable to Mm WT in Dd WT. This further corroborates previous evidence that Mm Δ*tesA* induces EsxA-dependent damage (Fig 3) and in absence of autophagy repair, more efficiently gains access to the cytosol, where it can grow without xenophagy restriction. Deletion of *tsg101* or *atg1* in Dd did not affect positively or negatively the strong attenuation of Mm ΔCEΔ*tesA* (Fig 4f).

To further confirm these conclusions, we quantified intracellular Plin-coating as a readout of cytosolic access (Fig 4g). As shown in Figure 3c, only Mm WT efficiently accessed the host cytosol in Dd WT. In contrast, the absence of autophagy-related repair in Dd Δ*atg1* cells increased Plin-coating for all Mm strains, and particularly for Mm Δ*tesA,* which increased from 13-26% to 54-63% of Plin-positive bacteria. This demonstrates that Mm Δ*tesA* in Dd Δ*atg1* reaches the host cytosol as efficiently as Mm WT. There is therefore a positive synergy between a Mm Δ*tesA* mutant weakened in its damaging capacity and a host deficient in repair.

Taken together, in absence of autophagy-mediated repair, EsxA is required and sufficient for cytosol escape, because deletion of *atg1* favours growth of both Mm WT and Mm Δ*tesA,* but not Mm ΔCE and Mm ΔCEΔ*tesA*. ESCRT and autophagy are therefore crucial to maintain MCV integrity following EsxA-induced membrane damage, while PDIM acts as a secondary virulence effector to overcome repair processes and allow progression to catastrophic damage and escape to the cytosol.

### The co-operative role of PDIM and EsxA is conserved in mammalian cells

To determine whether the sequential roles of EsxA and PDIM in membrane damage are conserved in animal phagocytes, we used murine microglial BV-2 cells, which have been successfully compared to the Dd-Mm model in previous studies (20, 64, 65). We first monitored the intracellular growth of the Mm mutants in BV-2 cells using automated high-content time-lapse microscopy and computer-assisted quantification of intracellular bacteria area as proxy for the bacteria load (Fig 5a-b, Fig S5). With the exception of Mm Δ*tesA*, all Mm mutants were strongly attenuated compared to Mm WT. The growth hierarchy observed in Dd was also conserved in BV-2 cells, with Mm Δ*tesA* ranking slightly below Mm WT, followed by Mm ΔRD1 and Mm ΔCE showing similar intracellular growth, and finally the two double Mm mutants displaying the strongest attenuation. Although the bacterial area of Mm Δ*tesA* was not significantly different from Mm WT, its statistical comparisons with other mutants were less pronounced than those obtained for Mm WT. Furthermore, the additive attenuation observed for Mm ΔRD1Δ*tesA* and Mm ΔCEΔ*tesA* suggests that the *tesA* deletion slightly attenuates the growth of Mm.

**Fig 5.**
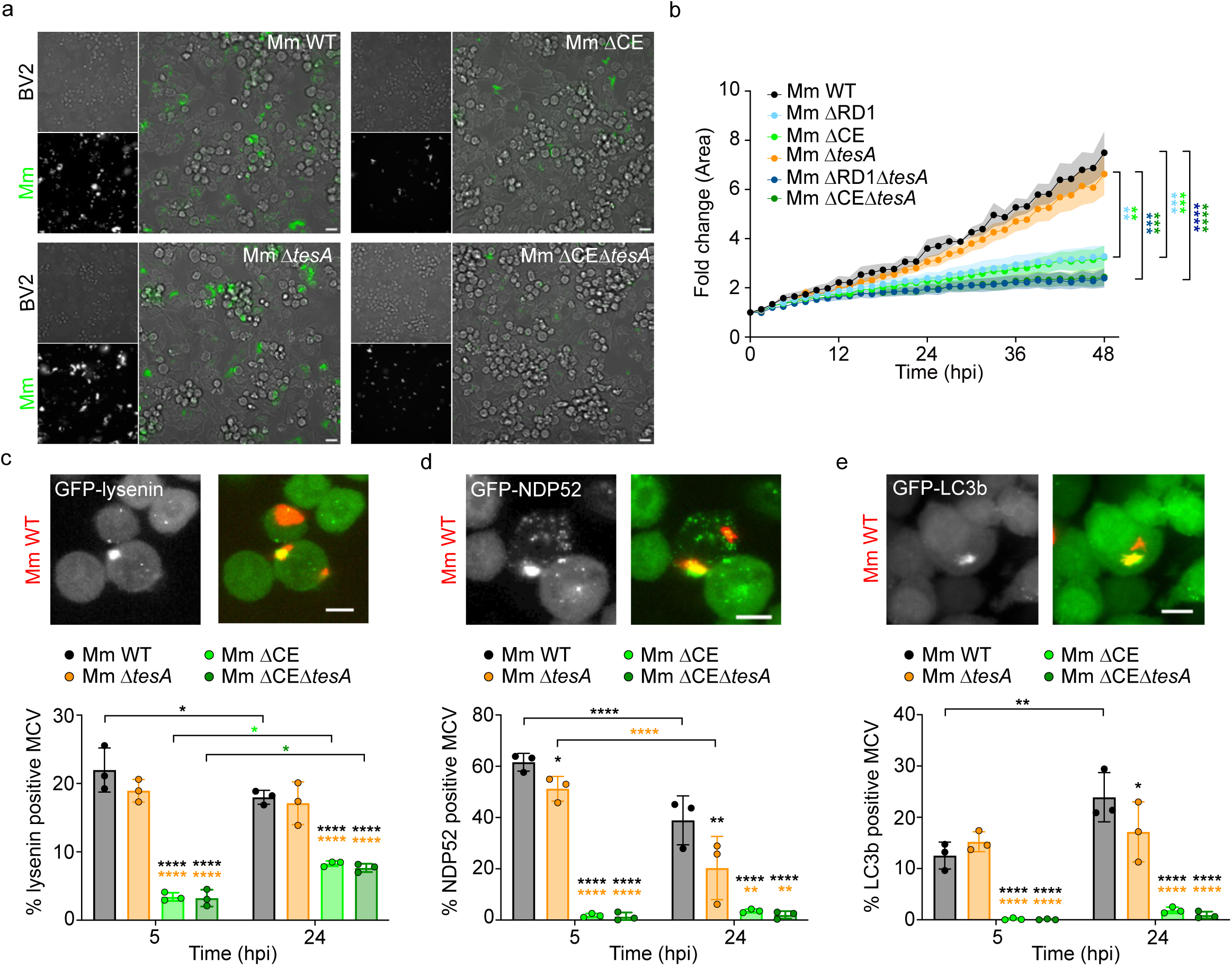
The sequential roles of EsxA and PDIM are conserved in mammalian cells. (a-b) BV-2 cells infected with GFP-expressing Mm mutants and monitored by high-throughput, high-content time lapse microscopy for 48 hours. (a) Representative pictures of infected cells at 48 hpi. Scale bar, 20 µm. (b) Growth curves of the Mm mutants measured as area sum of the GFP signal every 1.5 hpi and represented as the mean of fold change ± SEM (9 pictures per well, n = 4-6, N = 3-4, two-way ANOVA with Fisher’s LSD test). (c-e) BV-2 cells stably expressing GFP-lysenin (c), GFP-NDP52 (d) or GFP-LC3b (e) were infected with mCherry-expressing Mm mutants. After fixation at 5 and 24 hpi, the cells were imaged by high-throughput, high-content microscopy. The images are representative pictures of the different marker recruitments at the MCV of Mm WT. Scale bar, 10 µm. The graphs represent the average ± SEM of the automated quantification of marker recruitment at the bacteria (12 pictures per well, n = 3, N = 3, two-way ANOVA with Fisher’s LSD test).

Next, to assess the level of membrane damage induced by each mutant during BV-2 infection, we quantified the percentage of MCV reporter recruitment in cells stably expressing GFP-lysenin, GFP-NDP52 and GFP-LC3b at 5 and 24 hpi (Fig 5c-e). Lysenin is a toxin from the earthworm *Eisenia fetida* that specifically binds sphingomyelin (66). Recruitment of cytosolic GFP-lysenin to MCV membrane indicates the presence of small damage that leads to cytosolic exposure of lumenal sphingomyelin. As ESCRT recruitment during Dd infection, this reporter is then associated with early damage (Fig 5c). At an early time point, Mm WT and Mm Δ*tesA* showed similar percentages of GFP-lysenin positive MCVs, both significantly higher than observed for Mm ΔCE and Mm ΔCEΔ*tesA*. A similar recruitment profile was observed at 24 hpi, although there was a slight decrease in recruitment at Mm WT MCVs over time, and an increase at the MCVs of Mm ΔCE and Mm ΔCEΔ*tesA*. Thus, as observed in Dd, the absence of PDIM does not impair the ability of Mm to induce small membrane damage in BV-2 cells, whereas absence of EsxA does. The similar attenuations of Mm ΔCE and Mm ΔCEΔ*tesA* indicated that EsxA plays a predominant role in inducing repairable damage.

We next evaluated the association of GFP-NDP52, an autophagy adaptor, and GFP-LC3b, an autophagosomal membrane reporter, with the MCVs (Fig 5d-e). These reporters reflect the ability of Mm mutants to induce larger membrane damage. Mm WT showed 60% of MCVs positive for NDP52 at early time point, before decreasing to 40% at 24 hpi. Mm Δ*tesA* recruited slightly less NDP52 adaptor than Mm WT at 5 hpi, and this defect became more pronounced over time. Mm ΔCE and Mm ΔCEΔ*tesA* showed a strong recruitment defect with less than 5% of MCVs positive for GFP-NDP52 at all time points. In contrast, LC3b recruitment to Mm WT reached 12% at 5 hpi and increased to 24% at 24 hpi. Mm Δ*tesA* showed similar LC3b recruitment compared to Mm WT at 5 hpi but displayed a defect at 24 hpi. As observed with the autophagy adaptor, Mm ΔCE and Mm ΔCEΔ*tesA* showed almost no LC3b recruitment at their MCVs. These results indicated that all Mm mutants are impaired in inducing larger membrane damage, with defects observable starting from the autophagic adaptor recruitment to the stage of autophagosome formation.

Altogether, our results demonstrated that both EsxA and PDIM are required for optimal Mm growth in mammalian BV-2 cells. EsxA is required to induce both small sphingomyelin-positive wounds and larger autophagy repair-associated damage. In contrast, absence of PDIM does not affect Mm’s capacity to induce small sphingomyelin-positive damage but reduces the progression to larger membrane damage associated with the autophagy repair machinery. The similar attenuations observed for ΔCE and ΔCEΔ*tesA* across all marker recruitments indicate that EsxA plays a predominant role and is required for subsequent PDIM function. Thus, the sequential and synergistic action of EsxA and PDIM in inducing MCV membrane damage, initially characterized in Dd, is conserved in mammalian phagocytes.

## Discussion

In this study, we investigated how Mm regulates membrane damage/repair cycles to access the host cytosol in a timely and efficient manner, a pre-requisite for dissemination to naïve bystander cells. Our results reveal that EsxA and PDIM, although both known to participate in MCV membrane disruption, play distinct and non-equivalent roles during Mm infection (Fig 6). EsxA induces repairable membrane damage that triggers the complex recruitment of ESCRT and autophagy machineries. In contrast, PDIM is not required for the initiation of these repairable lesions but instead promotes their progression toward a catastrophic stage, ultimately enabling cytosol escape. Our data highlight a mechanism in which this progression does not simply reflect a gradual enlargement of a single damage but may involve a transition between distinct stages of membrane damage. Together, our findings support a refined model in which pathogenic mycobacteria tightly control membrane damage at the MCV, ensuring a sequential two-step process that balances host membrane repair with bacterial survival.

**Fig 6.**
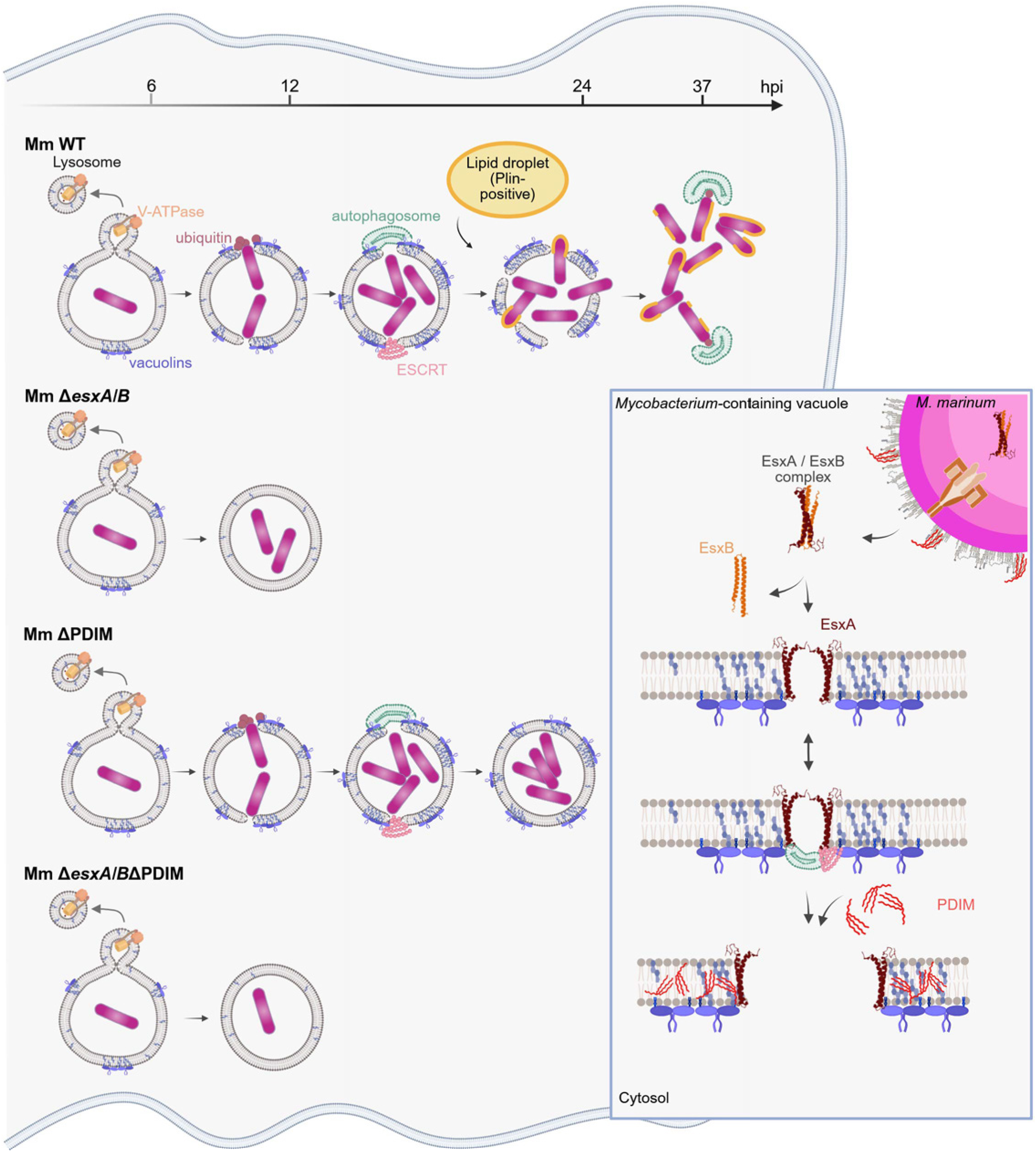
Model of the sequential roles of EsxA and PDIM during MCV membrane damage. Mm WT, once inside its MCV, induces small membrane damage through the secretion of EsxA by the ESX-1 system. These small damage recruit both the ESCRT and autophagy machineries for repair. PDIM then cooperates with EsxA to enlarge the lesions, preventing membrane repair and promoting bacteria access to the host cytosol. Mm Δ*esxA/B* mutant (Mm ΔCE) is rapidly constrained within its MCV as, in the absence of EsxA, no membrane damage occurs. Mm Δ*tesA*, unable to synthesise PDIM, displays an intermediate intracellular growth and level of damage. This Mm mutant induces small EsxA-dependent damage, which are repaired by ESCRT and autophagy, but remains trapped within its repaired MCV. The absence of both EsxA and PDIM in Mm leads to a stronger growth attenuation and absence of membrane damage, similarly to *esxA* deletion.

As a byproduct, our study also highlights the limitations of the classical haemolysis assay as *in vitro* membrane damage assay. While our infection-based results clearly identify a sequential damage process, the haemolysis assay detected only EsxA-dependent activity under our conditions. Similarly, Osman *et al*. reported their Mm PDIM-deficient mutant retained substantial haemolytic activity (54). Therefore, the haemolysis assay primarily identifies factors initiating membrane damage, but it is not well adapted to detect complex or sequential mechanisms. Culture conditions also appear to strongly influence the haemolysis results (67). Monitoring multiple damage and repair reporters during infection overcomes these limitations and should become a preferred approach to investigate membrane damaging factors.

The mechanisms of damage/repair cycles during Mtb/Mm infection have been extensively studied as they represent a critical step for bacterial survival in both macrophages and Dd. Both ESCRT and autophagy machineries are required to repair the damaged MCV (11, 13, 22, 24). MCV damage is often compared to sterile lysosomal damage induced by chemical agents such as LLOMe, because the two processes share similarities (68). While each repair pathway has been individually monitored by live imaging, fixed samples have so far been the only approach used to analyse them simultaneously (13). Successfully imaging both machineries in live cells revealed their high dynamics. A model in which these two pathways repair the damaged membrane sequentially, with ESCRT followed by autophagy, appears appropriate for the LLOMe damage/repair mechanism but overly simplistic for the MCV compared to our observations (13, 69). GFP-Vps32 and mCherry-Atg8A displayed a complex choreography with different patterns (dot, patch, full coverage), timings, and single or, most often, dual recruitment events. When both probes were present at the MCV, the reporters were mainly in juxtaposed, non-overlapping locations, as previously described (13). The higher prevalence of mCherry-Atg8A compared to GFP-Vps32 suggests that autophagy plays a predominant role in maintaining MCV integrity compared to ESCRT. Based on our quantitative analyses, ESCRT recruitment reflects early, repairable membrane lesions, whereas autophagy participates in initial responses to damage but is also strongly associated with extended damage that exceed ESCRT repair capacity. Previously in Dd, absence of the E3 ubiquitin ligase TrafE has been shown to lead to non-functional accumulation of ESCRT and autophagy reporters at damaged lysosomes (26), in contrast to the absence of the calcium sensor PefA which impairs Vps32 and Atg8A recruitment to damaged lysosomes and MCV (23). While PefA and TrafE coordinate ESCRT and autophagy, our observation that autophagy recruitment is reduced at the Mm Δ*tesA* MCV while ESCRT repair remains unchanged suggests that autophagy may also act in an ESCRT-independent manner in Dd, as demonstrated in mammalian cells (22, 27).

In mammalian cells, while canonical autophagy removes unrepairable lysosomes, ATG8ylation is emerging as a broader regulator of membrane repair pathways. Recent work indicates that ATG8ylation is a response to membrane damage triggered potentially independently of ESCRT but downstream of calcium leakage (31). ESCRT recruitment is proposed to occur downstream of ATG8ylation, to promote membrane repair in a context-dependent manner (32). ATG8ylation is also observed following MCV damage in macrophages (22, 30). In Dd, our observations are consistent with a form of ATG8ylation, rather than strictly canonical autophagy, contributing prominently to MCV damage/repair responses, as the MCV is not completely enclosed within an autophagosome (11, 13, 20, 23, 26).The roles of the different autophagy components should therefore be further dissected to clearly differentiate ATG8ylation from canonical autophagy and to distinguish ESCRT-dependent and -independent repair pathways.

Mycobacterial membrane-damaging factors have been widely explored. Although the initial membrane damage activity of EsxA was partially attributed to a minor detergent contamination (50), it is now well established that the ESX-1 effector EsxA confers to Mtb/Mm the ability to damage the MCV membrane (12, 33, 46, 59, 70). However, a controversy remains as one study attributes the damage activity to the ESX-1 system itself rather than to a specific substrate, reporting that direct contact between the bacterium and the host membrane is required (50). In our models, the similar intracellular growth attenuation of Mm ΔRD1 and Mm ΔCE provides evidence that the EsxA/EsxB dimer is the principal virulence factor secreted by the ESX-1 system. Our complementation experiment shows that very low levels of EsxA/EsxB are sufficient to restore the virulence in Dd, as previously demonstrated in mammalian cells (35). However, we did not rule out potential effects of EsxA/EsxB on the secretion of other ESX-1 substrates that could contribute to virulence. ESX-1 substrate secretion is tightly and hierarchically regulated, with the presence of EsxA/EsxB being essential for the secretion of other effectors (43). Recently, a pH-dependent switch in ESX-1 secretion has been described, suggesting an adaption of effector release to environmental conditions (67). Although *esxA/esxB* deletion is detrimental to the secretion of other ESX-1 effectors, the intrinsic membrane-interacting activity of EsxA supports a direct contribution to membrane disruption (20, 48, 49, 70). During infection, EsxA has been shown to specifically target sterol-rich host membrane microdomains in both Dd and mammalian cells (20). Whether EsxA can be considered a classical sterol-dependent bacterial pore-forming toxin remains debated. In addition to its sterol requirement similar to those pore-forming toxins, EsxA induces NINJ1 oligomerization and plasma membrane rupture as demonstrated for perfringolysin O from *Clostridium perfringens* (71). Recently, EsxA has been proposed to rupture membranes through fibrillation within compartments, inducing vesiculation and membrane remodelling (52). Although the precise mechanism of EsxA-mediated membrane damage remains unclear, we confirmed here that EsxA induces MCV membrane damage, leading to recruitment of both ESCRT and autophagy-related repair machineries.

Initially, PDIM was shown to participate in phagocytosis by modifying the lipid organization at the plasma membrane of macrophages to facilitate Mtb entry (72). More recently, PDIM has also been characterized as a direct membrane damaging factor. These studies identified membrane damage defects for Mtb or Mm strains carrying mutations in the PDIM synthesis pathway, based on reduced recruitment of ubiquitin, galectin 3/8 and/or autophagy reporters, as well as impaired cytosolic access measured using a cell wall-associated β-lactamase assay (53-56). Our results confirm and extend these observations, showing a significant reduction in autophagy reporter recruitment in both Dd and BV-2 models, and an absence of cytosolic access in Dd for the Mm Δ*tesA* mutant. Overall, these reporters mainly reflect extended damage rather than newly induced repairable lesions. In contrast, both ESCRT subunits and lysenin were efficiently recruited to the MCVs of Mm mutants unable to produce PDIM. This uncoupling between repairable and catastrophic damage indicates that PDIM is not required for damage initiation but specifically for the transition toward catastrophic, non-repairable damage, ultimately allowing full access to the cytosol. Mm ΔCEΔ*tesA* recapitulates all reporter recruitments observed for Mm ΔCE, demonstrating that EsxA function is predominant over that of PDIM. Cooperation between EsxA and PDIM has already been observed in the uptake-independent killing of macrophages by Mtb aggregates (73). For membrane damage, Augenstreich *et al*. hypothesized that PDIM potentiates EsxA activity (56) and corroborated this *in vitro* using liposomes (59). Our model is also consistent with the study from Bah et al., demonstrating that PDIM promotes autophagy-dependent damage in the presence of a functional ESX-1 system (74). While studying the mechanism of Mtb PDIM-dependent entry into macrophages, PDIM was shown to target cholesterol (72). Cholesterol is also essential for PDIM function during zebrafish infection (58). PDIM is transferred from the Mm or Mtb envelope to macrophage MCV and other membranes where it adopts a conical shape that destabilizes membranes (58, 75, 76). As EsxA requires sterol-rich microdomains to damage the MCV (20), our data support the hypothesis that PDIM enhances EsxA activity during infection. Here, we provide evidence that Mm employs a two-step mechanism to damage the MCV membrane: EsxA initiates the damage process, while PDIM is required for progression toward a catastrophic state, potentially by promoting clustering of sterols in the MCV membrane, thereby facilitating further EsxA-mediated membrane damage. An open question remains concerning the precise PDIM mode of action, specifically whether PDIM function requires shedding from the mycobacterial cell wall, as previously reported (58, 75), or direct contact between the bacterium and the host membrane.

Distinctively, the time necessary for Mtb/Mm cytosol access is considerably longer than that reported for other intracellular bacteria. *Shigella flexneri* or *Listeria monocytogenes*, two of the best characterized bacterial pathogens that reach the host cytosol, require only minutes to rupture their vacuole (77, 78). In contrast, the delayed cytosolic escape of Mtb/Mm is likely due to the damage/repair cycles that do not occur for these other bacteria. This late vacuolar escape may be critical for some metabolic adaptations and/or to overcome the host antimicrobial defence strategies. Supporting this hypothesis, a transposon sequencing analysis of Mm following Dd infection revealed a fitness defect for Mm mutants carrying insertions in genes involved in ESX1 secretion or PDIM synthesis, similarly to mutants deficient in metabolic enzymes (64). Interestingly, transposon insertions disrupting enzymes specific to PGL synthesis conferred an intracellular fitness advantage to Mm. This phenotype was interpreted as an indirect gain of function effect resulting from metabolic rewiring toward the PDIM pathway, boosting PDIM synthesis and consequently Mm virulence (64).

It is also important to note that an Mm mutant deficient in PDIM/PGL synthesis exhibits altered ESX-1 secretion (79, 80). While the phenotypes observed in our study appear to be linked to PDIM/EsxA cooperation in membrane damage, we cannot exclude that altered ESX-1 secretion contributes to these effects. The additive effect observed for the double Mm mutants in intracellular growth assays could reflect this secretion defect. Indeed, as PDIM damage activity requires EsxA, an intracellular growth similar to Mm ΔCE would have been expected for Mm ΔCEΔ*tesA*. Moreover, despite strong reductions, damage reporter recruitments and cytosolic access never reach a zero level with the different Mm mutants tested, suggesting that additional bacterial factors may contribute to membrane damage. For example, it was suggested that the ESX-1 effector EspB can replace EsxA/EsxB during uptake-independent killing in macrophages, indicating potentially conserved roles between these effectors (73). A study using Mm transposon mutants targeting ESX-1 genes identified strains that were unable to secrete EsxA but nevertheless damaged their MCV and accessed the host cytosol (51). In contrast, Mm Δ*espE* and Mm Δ*espF* are able to secrete EsxA/EsxB but show reduced haemolytic activity and attenuated intracellular growth in macrophages (81). EspA and EspC may also compensate for EspE and EspF function under acidic conditions (67). Strikingly, deletion of the fatty acyl-CoA ligase FacL6 from Mm alters the ESX-1 gene expression profile, likely leading to exacerbated MCV damage (82). Beyond ESX-1, additional factors such as the Mm phosphatases and the Mtb sphingomyelinase SpmT have also been linked to MCV lysis and membrane damage and represent potential candidate regulators for further investigation, particularly in combination with EsxA and PDIM (83, 84).

The importance of a sequential mechanism between EsxA and PDIM during *in vivo* infection remains uncertain. Mm Δ*tesA* was previously shown to be attenuated in the zebrafish model. Although the Mm Δ*tesA* mutant was also attenuated to a lesser extent than ΔRD1 in their Dd model, it exhibited a similar zebrafish survival phenotype as ΔRD1 (61). Moreover, an Mtb strain unable to produce PDIM is also less virulent in mice. This attenuation has been linked to the ability of PDIM to inhibit LC3-associated phagocytosis and autophagy rather than to membrane damage (85).

Using host cells with different genetic backgrounds can also help to characterize the damage activity of Mtb/Mm mutants. As demonstrated here with the autophagy deficient Dd and Mm Δ*tesA*, a weakened host can partially rescue a weakened bacteria mutant. Notably, the restoration of cytosol access and growth for Mm Δ*tesA* in Dd Δ*atg1* indicates that PDIM is specifically required to overcome host autophagy-related repair mechanisms instead of initiating membrane damage. However, it is important to note that deletion of a specific repair pathway does not eliminate all host defences, for example, xenophagy may remain functional. Consequently, after premature vacuole escape, bacterial growth may still be restricted, as observed in ESCRT-deficient Dd (13). Depending on the bacterial growth attenuation monitored in WT cells, it might be difficult to observe a further attenuation as seen here for Mm Δ*tesA* in Dd Δ*tsg101*.

Overall, our work aligns with the hypothesis proposed in recent years and extends our understanding of the cellular processes involved, allowing us to propose a refined model of MCV membrane damage (Fig 6). EsxA, once secreted into the MCV through the ESX-1 system, induces small membrane lesions that trigger the recruitment of ESCRT and autophagy repair machineries. PDIM subsequently promotes the progression of these lesions toward a catastrophic state, releasing the bacteria into the host cytosol. This sequential model is observed in both amoeba and mammalian cells, highlighting the evolutionary conservation of membrane damage/repair processes.

## Materials & Methods

### Cell lines

The Dd strains used in this study are listed in Table S1. Cells were axenically grown in 10 cm Petri dishes at 22°C in HL5 or HL5c medium (Formedium) supplemented with 100 U/mL of penicillin and 100 µg/mL of streptomycin (Invitrogen).

Primers and engineered plasmids to generate the Dd cell lines expressing fluorescent proteins are listed in Tables S2 and S3. Plasmids were transfected into Dd by electroporation following the protocol from Perret *et al.*, 2026 (62). Hygromycin and G418 were used for selection at a concentration of 50 and 7 µg.mL^-1^, respectively.

Mammalian cell lines used in this study are listed in Table S1. Cells were cultured at 37°C with 5% CO_2_ in DMEM medium (Gibco) supplemented with 10% heat inactivated fetal bovine serum (Invitrogen) and 100 U/mL of penicillin and 100 µg/mL of streptomycin (Invitrogen).

Plasmids to generate stable murine microglial BV-2 cell lines expressing fluorescent proteins are listed in Tables S3. Retrovirus were produced in HEK293T cells after cotransfection of the plasmid containing the DNA of interest with the packing vectors pMD-VSVG and pMD-OG, each being a gift from Felix Randow. The retrovirus were then used to infect BV-2 cells and cells stably expressing the genes of interest were selected with blasticidin at a concentration of 5 µg.mL^-1^.

### Bacterial strains

The Mm M strains used in this study are listed in Table S1. The bacteria were cultured on 7H10 or 7H11 agar plates (Difco) supplemented with 0.5% glycerol, 10% OADC (Becton Dickinson) and in 7H9 (Difco) supplemented with 0.2% glycerol, 10% OADC or ADC and 0.05% Tween 80 (Sigma Aldrich) or tyloxapol (Sigma Aldrich). They were grown at 32°C and at 150 rpm for the liquid cultures in presence of 5 mm glass beads to minimize bacterial aggregation. *M. smegmatis* mc^2^155 was used in the phage transduction process and grown aerobically at 37°C in the same medium than Mm but without shaking to reduce aggregation.

Primers and engineered plasmids are listed in Tables S2 and S3. The selection of Mm mutants deleted for the *tesA* gene was realized with hygromycin 50 µg.mL^-1^. The complemented strains were produced by electroporation of the appropriate complementation plasmid in the corresponding Mm mutants, and the genomic integration of the plasmid were selected and then maintained with apramycin 20 µg.mL^-1^. The strains with the fluorescent plasmids were selected and maintained with hygromycin 50 µg.mL^-1^ and/or kanamycin 20 µg.mL^-1^.

*Escherichia coli* Top10, DH5α and HB101 were used for cloning and grown at 37°C in LB medium. Hygromycin (100 µg.mL^-1^), kanamycin (50 µg.mL^-1^) and apramycin (30 µg.mL^-1^) were added to the media, if required.

### DNA constructs

All Primers and engineered plasmids used in this work are listed in Tables S2 and S3, respectively. Q5 (NEB), phusion (NEB) or PrimeStar Max (Takara) polymerases were used for PCR amplifications. The classic restriction enzyme strategy was used to construct all the plasmids, with calf intestinal alkaline phosphatase (NEB) to prevent re-ligation of linearized plasmids and T4 ligase (NEB) to ligate DNA fragments. *E. coli* HB101 was used to clone allelic exchange substrates into phAE159 phage DNA while *E .coli* Top10 and DH5α were used for the other cloning. Each PCR product and plasmid sequences were validated by sequencing.

### Transformation of plasmids into mycobacteria

Electrocompetent mycobacteria were prepared as in the established protocols (86). In short, a 10 mL bacterial culture was grown to an OD_600_ of 0.8 to 1, washed at least twice with 10 % glycerol containing 0.05 % tyloxapol and finally concentrated to 1 mL. For *M. smegmatis* or Mm, the procedure was performed on ice with ice-cold glycerol/tyloxapol or at room temperature (RT), respectively. 1-5 µg of plasmid DNA was electroporated into 400 µl electrocompetent mycobacteria using electroporation cuvettes with 0.2 cm electrode gap. A BioRad Gene Pulser was set to 2.5 kV, 1 kΩ and 25 µF capacitance. 1 mL of 7H9 broth was added for recovering after electroporation. *M. smegmatis* was recovered for 2 hours at 37°C without shaking, whereas Mm was incubated 4 hours up to overnight at 30-32°C before plating on 7H10 or 7H11 plates containing the appropriate antibiotics.

### Generation of defined gene deletion mutants in Mm by transduction

The allelic exchange substrates (AES) for the gene *tesA* needed for specialized phage transduction were generated as described in detail previously (83). The AES are designed to replace the genes of interest in Mm M with a *γδres*-*sacB*-*hyg*-*γδres* cassette comprising the *sacB* and a hygromycin resistance gene flanked by *res* sites of the γδ-resolvase. The flanking regions of *tesA* were amplified using the primer pairs oHK47/oHK48 and oHK49/oHK50. The PCR products were digested with DraIII or Van91I, respectively, and ligated with Van91I-digested p0004S vector fragments to gain the plasmid pHK12. The DNA of the temperature-sensitive phage phAE159 and pHK12 were linearized with PacI before ligation to gain pHK23. High-titer phages of phAE159Δ*tesA* were prepared in *M. smegmatis* to perform specialized transduction in Mm to yield Δ*tesA*. The mutants obtained were verified by PCR using primer pairs with one primer located in the genome (Table S3: oHK96/97) and the other primer located in the resistance cassette (Table S3: oHK33 and oHK36) and sequencing of the amplicon.

### Generation of defined gene deletion mutants in Mm by suicide vector

The Mm Δ*pks15/1* mutant was generated using the suicide vector strategy published by Arnold *et al.* (87). Briefly, the upstream and downstream region of the gene *pks15/1* were amplified and were then both ligated into the pINIT vector thanks to their specific and oriented BspQI restriction sites. The corresponding pINIT-*pks15/1* plasmid was verified by sequencing. The DNA fragment containing the upstream and downstream *pks15/1* regions was transferred to the pKOΔ*galK* plasmid (addgene #110090). The new pKO-*pks15/1* plasmid was treated by UV light (100 000 µj/cm2) before 5 µg was transformed into Mm M strain. The mutants obtained were verified by PCR using primer pairs with one primer located into the genome and the other into the plasmid.

### Secretion assay

50 mL mycobacteria cultures were grown to an OD_600_ of 0.8-1, washed twice with PBS 1x, and the quantity of bacteria for new cultures of 50 mL at OD_600_ initial 0.8 was incubated at 32°C in Sauton medium for 48 hours. Bacteria and supernatants were separated by centrifugation at 3 000 rpm 10 min. Bacteria pellets were washed with TBS 1x, before being resuspended in TBS 1x with cOmplete EDTA-free protease inhibitors (Roche), sonicated and lysed by bead beating (3 cycles of 30 s with 15 s break interval). Bacterial lysates were centrifuged at 13 000 rpm for 5 min 4°C to recover the soluble fractions. Supernatants were filtrated through 0.22 µm filter to eliminate the remaining bacteria. cOmplete EDTA-free protease inhibitor is added to each supernatants before concentrating the samples with Amicon Ultra-4 3K columns to obtain the culture filtrates. Protein concentrations of cultures filtrates and bacterial lysates were determined by Nanodrop and 650 µg total proteins from bacterial lysates or 75 µg total proteins from culture filtrates were analysed by western blot. 4-20% Bis-Tris mPAGE gels (Millipore) were used to separate the proteins, before transfer on 0.2 µm nitrocellulose membrane (Amersham). The proteins of interest were detected with the following primary antibodies: anti-ESAT6 (BEI; NR-13803; 1/5 000), anti-CFP10 (BEI; NR-13801; 1/10 000); anti-GroEL (Enzo Life Sciences; ADI-SPS-875; 1/10 000); anti-Mpt32 (BEI; NR-13807; 1/50 000).

### Lipid extraction and thin-layer chromatography to visualize PDIM and PGL

40 mL mycobacteria cultures were grown to an OD_600_ of 0.8-1 in 7H9 (Difco) supplemented with 0.2% glycerol, 10% ADC and 0.05% tyloxapol (Sigma Aldrich). The bacterial pellets were washed once in PBS 1x, resuspended in ddH_2_O and incubated in CHCl_3_:MeOH (1:2) for 24 h at 150 rpm in the dark. After centrifugation, the supernatants were mixed to CHCl_3_: ddH_2_O (1:1) and then the organic phases were again mixed to CHCl_3_: ddH_2_O (1:1) before being washed a last time with ddH_2_O. Nitrogen dried extracted lipids were resuspended in CHCl_3_ in adapted volume to normalize the extracts to their total protein concentration. Running system used to visualize PDIM was petroleum ether:diethylether (90:10) while the PGL was migrated in chloroform:methanol (95:5). The different lipids were revealed by a charring solution containing sulfuric acid, MnCl2 and methanol.

### Haemolysis assay

20 mL mycobacteria cultures were grown to an OD_600_ of 0.8-1, washed twice with DPBS 1x, and 1.4*10^8^ bacteria were centrifuged with washed defibrinated sheep red blood cells (RBC - Thermo scientific). The pellets mycobacteria/RBC were incubated 2 h at 32°C before resuspension and centrifugation. The OD_405_ of the supernatants were determined using a SpectraMax i3 (Molecular devices). The percentage of haemolysis was calculated as below and normalized to the haemolysis obtained with Mm WT.

% haemolysis = (OD_405_ sample- OD_405_ DPBS) / (OD_405_ 0.1% Triton X100- OD_405_ DPBS)

### Dd cytosol-membrane separation

The cellular fractionations were performed as previously described in Perret *et al.*, 2026 (20). Briefly, 10^9^ cells were washed in Sorensen-Sorbitol before being resuspended in HESES buffer (20 mM HEPES, 250 mM Sucrose, 5 mM MgCl2, 5 mM ATP) with cOmplete EDTA-free protease inhibitors (Roche). Cells were broken using a ball homogenizer with a 10 µm clearance and the dead cells and nuclei were eliminated by centrifugation. The post-nuclear supernatant was diluted in HESES buffer and centrifuged at 35 000 rpm in a Sw60 Ti rotor (Beckman) for 1 h at 4°C to recover the cytosol (SN, supernatant) and the membranes (MB, pellet). The protein concentration of each extract was determined by Bradford assay.

### Plin-binding assay

20 mL mycobacteria cultures were grown to an OD_600_ of 0.8-1, washed twice with PBS 1x, and 1*10^6^ bacteria were incubated with 550 µg total cytosolic proteins for 1 h at RT under rotation. The samples were then divided into two. Sodium azide 5 mM were added to half of the sample, and these samples were analysed by FACS using a FACS cell sorter SONY SH800. The other half of the samples were transfer on poly-lysine coated coverslips, fixed with paraformaldehyde 4 %, rinsed with PBS 1x- glycine 100 mM and mounted with Prolong gold antifade mounting medium (Invitrogen). The samples were imaged using 63×1.4 NA oil immersion objective on a Leica SP8 confocal microscope.

### *In vitro* Mm growth

5*10^5^ bacteria from fresh cultures were seeded in 7H12 medium supplemented with OADC and tyloxapol in triplicate in 96 black wells plate with transparent bottom. The fluorescence from the bacteria was measured every hour for 48 h at 32°C using a SpectraMax i3 (Molecular devices). The doubling times were calculated from the obtained growth curves.

### Phagocytosis assay

Phagocytosis assays were made as previously described in Sattler *et al.* (88). Briefly, 10^7^ Dd cells were resuspended and incubated 2 h under agitation (150 rpm) in 6 well plates. 24 h Mm cultures at an OD_600_ of 0.8-1 were concentrated to obtain a 500 µl bacteria sample with 10^9^ bacteria (MOI 100). Bacteria were syringed to eliminated clumps and were added to the cells under agitation after having collected the sample T0. At the mentioned time point, 1*10^6^ cells were collected, and the phagocytosis process was stopped with 5 mM sodium azide before acquiring the data using a FACS Gallios.

### Infection

Infections were performed as previously described (89) with few modifications. 24 h prior the experiment, 1.3*10^7^ Dd or 5*10^6^ BV-2 cells were seeded in dishes without antibiotic and Mm strains were cultured to achieve an OD_600_ of 0.8-1. Bacteria were then rinsed and syringed before their addition to the cells at a MOI of 10 for Mm WT, Mm Δ*tesA* and Mm Δ*pks15/1* or 20 for Mm ΔRD1, Mm ΔCE, Mm ΔRD1Δ*tesA* and Mm ΔCEΔ*tesA*. Infections were synchronized by centrifugation, and 20 min phagocytosis were then allowed before washing the extracellular bacteria. The infected cells were resuspended in medium with bacteriostatic dose of 5 µg/mL streptomycin and 5 U/mL penicillin to prevent bacterial extracellular growth, and the appropriate quantity of cell was seeded in the relevant plate format.

To follow the intracellular growth in Dd by plate reader, 5*10^4^, 1*10^5^ or 1.5*10^5^ cells were plated in black 96 wells plate with flat transparent bottom. The plates were sealed with a gaz permeable membrane and the green fluorescence was recorded for 72 h with a Agilent BioTek Synergy H1 plate reader. For the High-content experiments, 7.5*10^3^ Dd for the intracellular growth or 1.2*10^4^ and 5*10^3^ Dd for the recruitment were plated per well in triplicate in IBIDI 96 wells plate and the plate were sealed. The infected cells were imaged using a 40x or a 60x water immersion objective on a Molecular Devices ImageXpress Micro Confocal microscope. The infected double reporter Dd cell line was settled in 4 wells IBIDI slides with 2*10^5^ cells per well, immobilized with an agarose sheet on the top and imaged using either a Nikon Ti / CSU-W1 spinning disc Confocal microscope with a 100x/1.49NA oil objective or a Leica Stellaris 8 Falcon microscope equipped by a 63x/1.4NA oil immersion objective. 4*10^4^ or 1*10^5^ infected BV-2 cells were seeded in triplicate in 96 wells IBIDI plate for intracellular growth or marker recruitment, respectively. Once sealed or fixed, the plates were imaged using a 40x water immersion objective on a Molecular Devices ImageXpress Micro Confocal microscope.

### Analysis

The data acquired by FACS were analysed with the online software Floreada (Floreada.io). The pictures and movies acquired by confocal microscopy were evaluated using ImageJ/Fiji. All the data obtained from the HC microscope were process using the MetaExpress software to perform automated single-cell analysis after segmentation (62, 89). The recruitment data were then imported to a R pipeline to extract the percentage of recruitment per MCV. The GraphPad Prism was used to perform statistical analysis and to plot graphs. The sample sizes are presented in the appropriate figure legends (n = number of technical replicates; N = number of biological replicates). The level of significance is indicated as \**P* < 0.05; \*\**P* < 0.01; \*\*\**P* < 0.005, \*\*\*\**P* < 0.0001.

## Acknowledgements

We are thankful to F. Randow (MRC-LMB Cambridge) for providing the retroviral plasmids: GFP-lysenin, GFP-NDP52 and GFP-LC3b. We acknowledge the staff of the Bioimaging Center photonic and ACCESS Geneva facility at the faculty of Sciences of the University of Geneva for their help. The following reagent was obtained through BEI Resources, NIAID, NIH: Polyclonal Anti-*Mycobacterium tuberculosis* ESAT6 (Gene Rv3875) (antiserum, Rabbit), NR-13803; Polyclonal Anti-*Mycobacterium tuberculosis* CFP10 (Gene Rv3874) (antiserum, Rabbit), NR-13801; Polyclonal Anti-*Mycobacterium tuberculosis* Mpt32 (Gene Rv1860) (antiserum, Rabbit), NR-13807.

This work was supported by the Swiss National Science Foundation grants 310030_188813 and 310030_219364 (TS) as well as the SystemsX.ch project HostPathX (HH, TS).

## Figure legends

**Fig S1.**
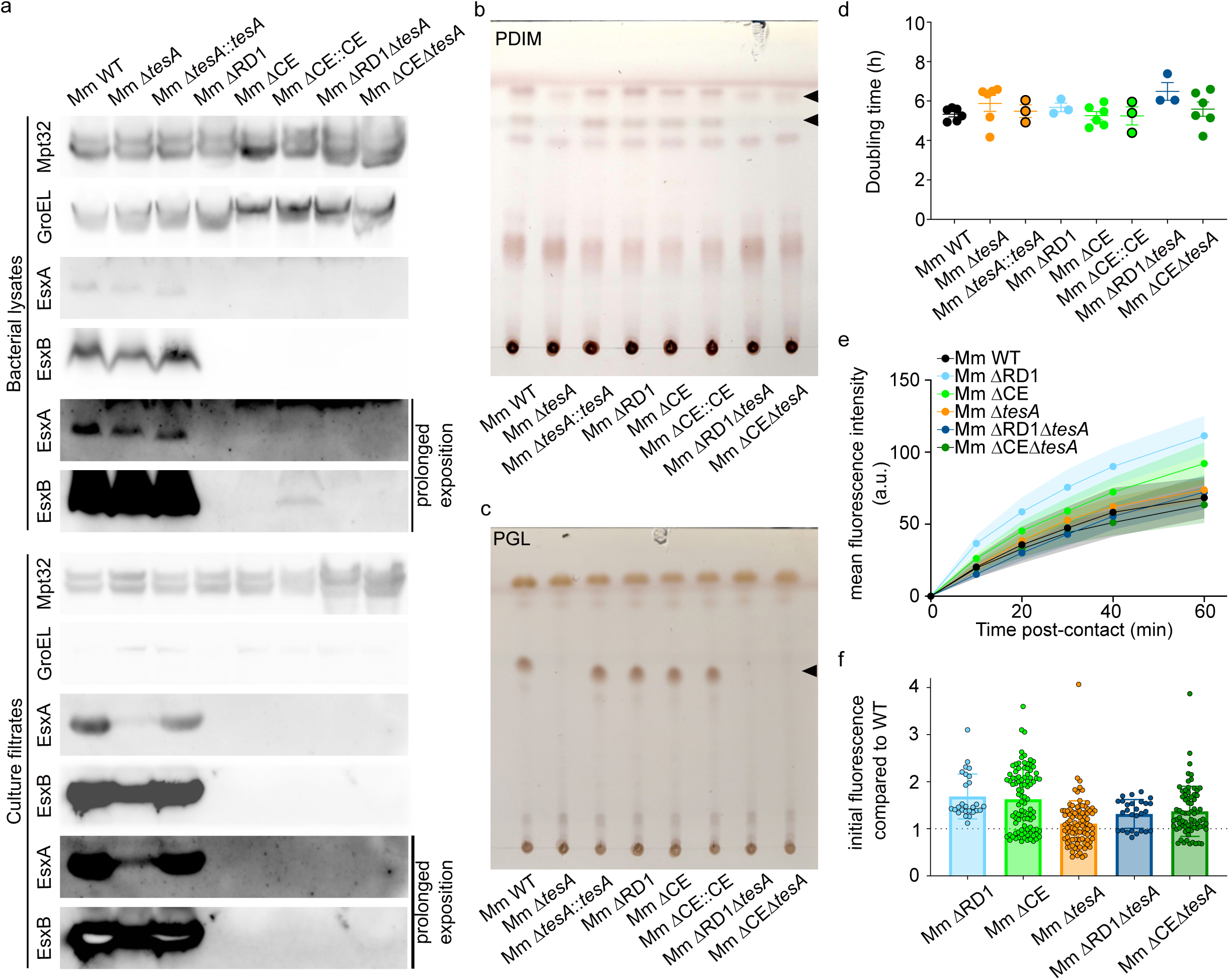
Mm mutants present neither an *in vitro* growth defect nor an uptake defect. (a) Western blot on Mm lysates (top) or Mm secreted proteins (bottom) performed after secretion assay on the set of Mm mutants. Mpt32 corresponds to a loading control for the secreted fractions. GroEL is a control for lysis. The blots are representative of three independent experiments. (b-c) Thin layer chromatography of total lipid extracts from the set of Mm mutants. The plate (b) migrated in petroleum ether: ethyl acetate (98:2) to visualize PDIM and the plate (c) in chloroform: methanol (96:4) to reveal PGL. The pictures are representative of three independent experiments. (d) *In vitro* doubling time of the Mm mutants calculated from *in vitro* growth assay. GFP-expressing Mm mutants were cultivated in 7H12 medium and the green fluorescence intensity was monitored for 48 hours by plate reader. Each data point corresponds to one biological replicate which is the average of technical triplicate (n = 3, N = 3-6, one-way ANOVA with Dunnett’s multiple comparison to Mm WT). (e) Phagocytosis assay determining the uptake ability of each Mm mutants. Dd were shaken with GFP-expressing bacteria and the green fluorescence was measured by flow cytometry at the indicated time points. The graph shows the average of mean fluorescence intensity ± SEM (n = 1, N = 3, two-way ANOVA with Dunnett’s multiple comparisons to Mm WT). a. u., arbitrary unit. (f) Initial fluorescence data obtained from each plate reader experiments to measure the intracellular growth of the Mm mutants, normalized to Mm WT. Each data point represents the initial fluorescence of one well from 3-10 independent experiments performed in technical triplicates.

**Fig S2.**
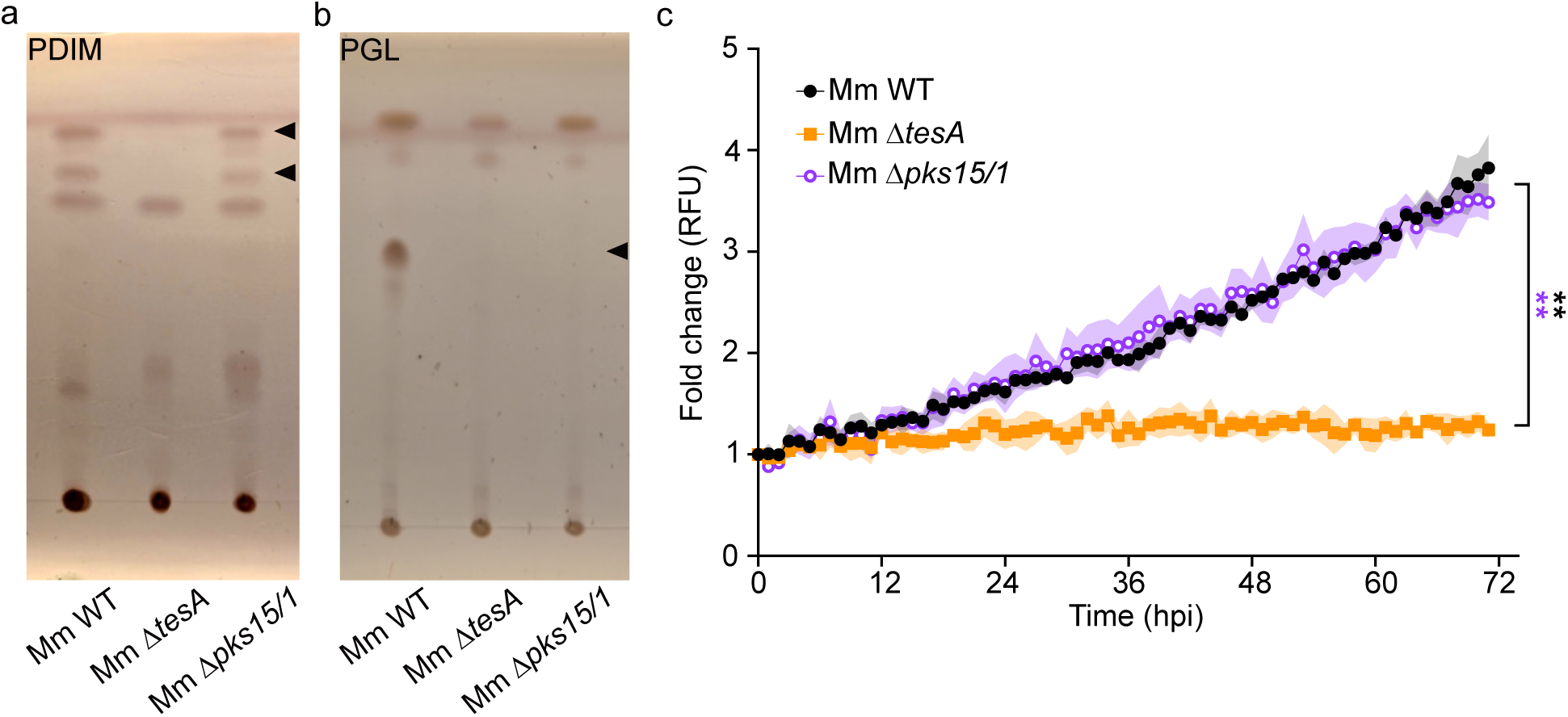
The absence of PGL does not influence the intracellular growth of Mm. (a-b) Thin layer chromatography of total lipid extracts from Mm WT, Mm Δ*tesA* and Mm Δ*pks15/1* mutants. The samples in (a) migrated in petroleum ether: ethyl acetate (98:2) to visualize PDIM while the plate in (b) was incubated in chloroform: methanol (96:4) to reveal PGL. The pictures of the plate are representative of three independent experiments. (c) Dd were infected with GFP-expressing Mm mutants and green fluorescence intensity was monitored for 72 hours by plate reader. The graph presents the average fold change ± SEM (n = 3, N = 3, two-way ANOVA with Fisher’s LSD test). RFU, relative fluorescence unit.

**Fig S3.**
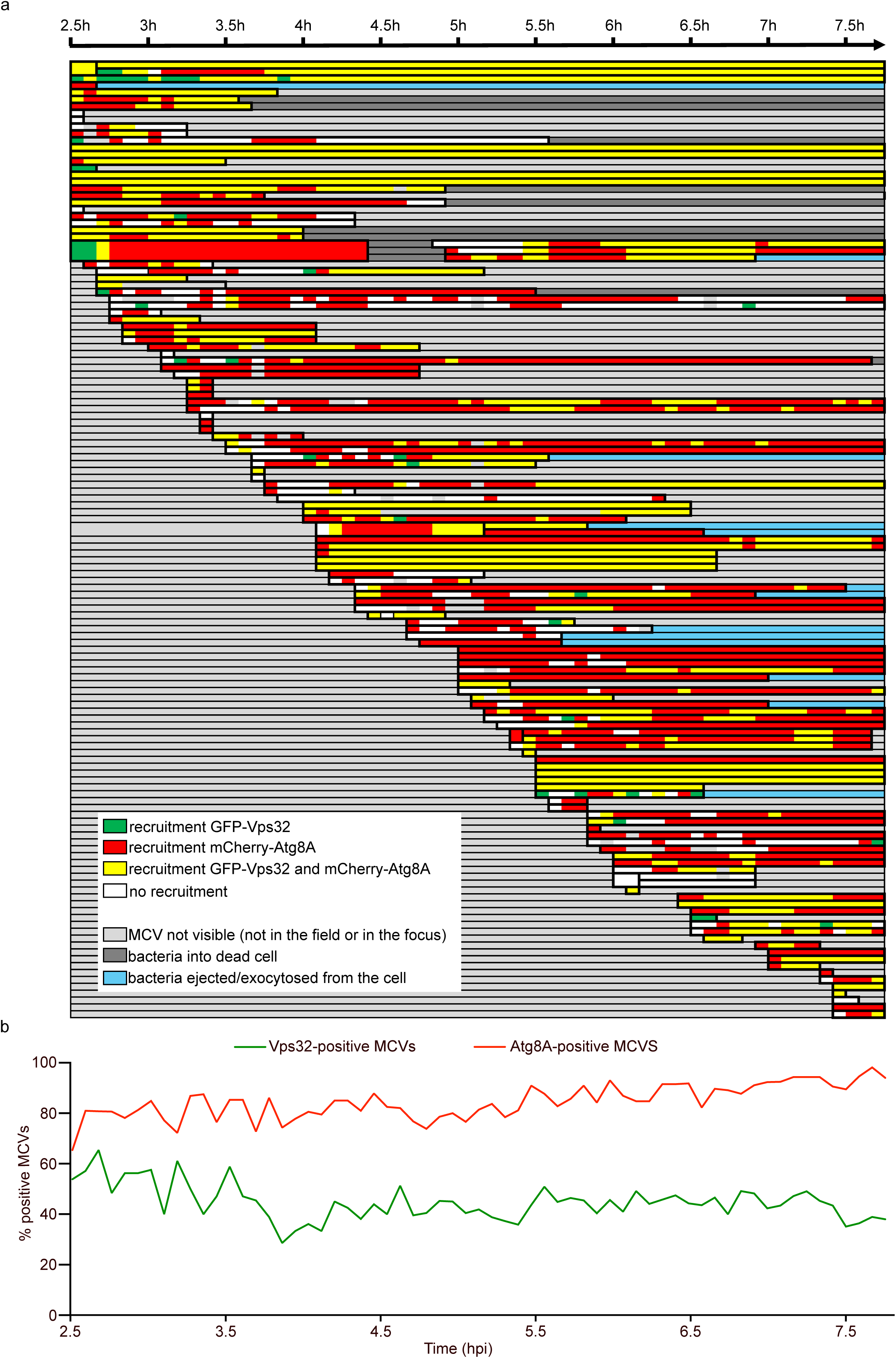
ESCRT and autophagy are differently but dynamically recruited at the MCV. (a-b) Dd expressing both GFP-Vps32 and mCherry-Atg8A were infected with EBFP-expressing Mm WT and monitored by live confocal imaging for several hours post-infection with pictures taken every 5 minutes. (a) Schematic representation of the reporter recruitment to MCVs tracked through time. Manual tracking of 140 independent MCVs, each line represents one MCV. (b) The graph represents the average recruitment of both reporters calculated from the independent MCV tracking above.

**Fig S4.**
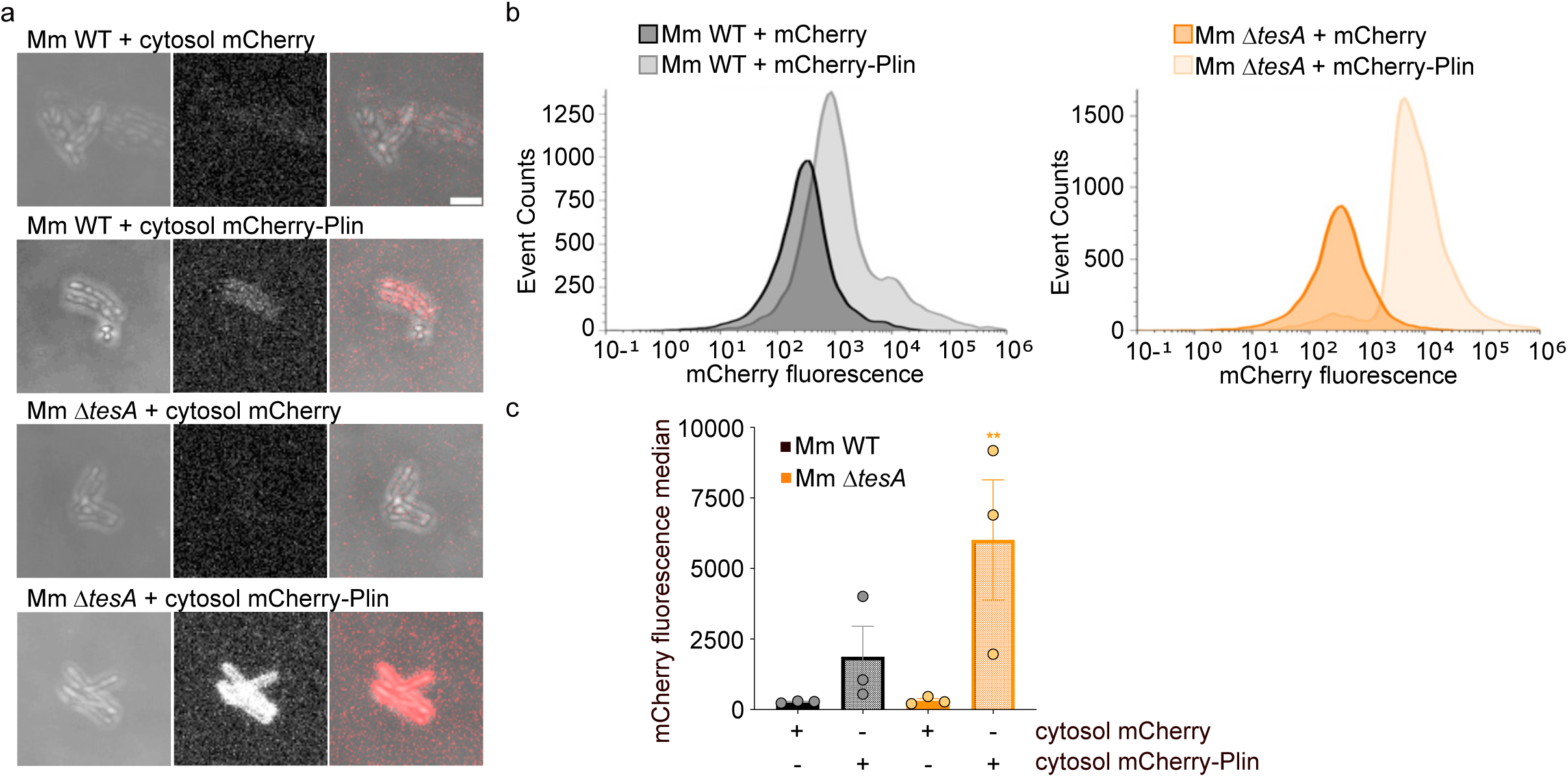
Mm Δ*tesA* mutant binds Plin proteins *in vitro*. (a-c) Plin-binding assay to determine the level of perilipin binding to Mm *in vitro*. (a) The images are representative pictures of Mm WT and Mm Δ*tesA* incubated in cytosols extracted from Dd expressing mCherry or mCherry-Plin. Scale bar, 2 µm. (b) and (c) are FACS analysis of the Plin-binding assay. (b) show the mCherry fluorescence intensity depending on the event count from one representative experiment. (c) displays the mCherry fluorescence median of each sample from independent biological replicate. (N = 3, one-way ANOVA with Fisher’s LSD test).

**Fig S5.**
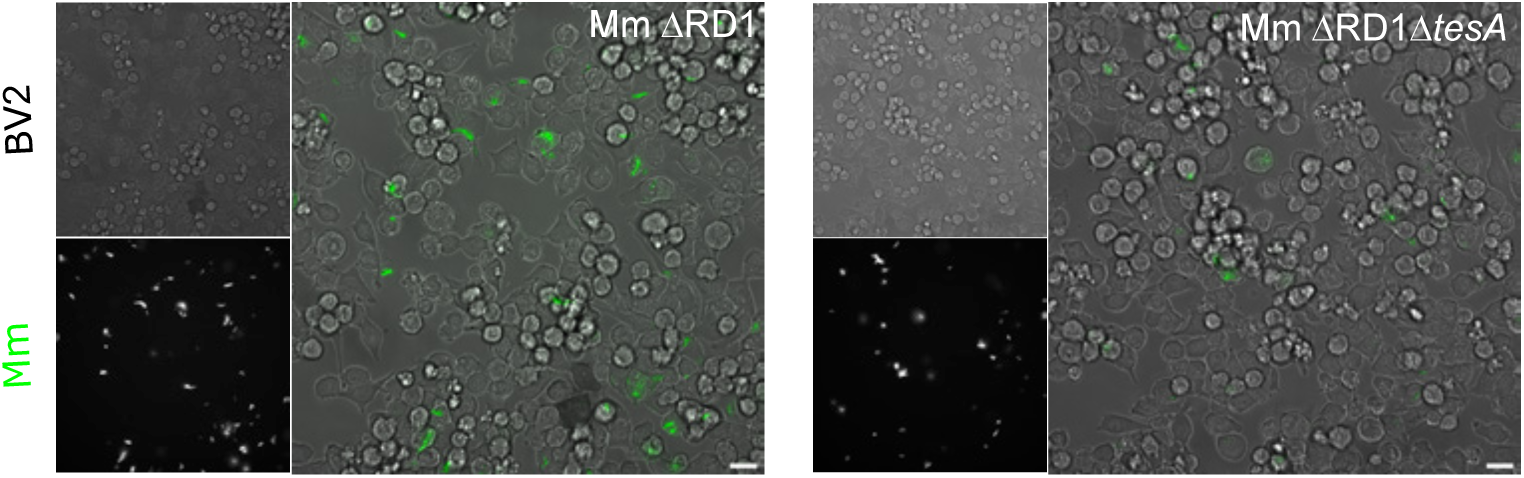
Mm ΔRD1 and Mm ΔRD1Δ*tesA* are attenuated in BV-2 cells. Representative pictures of BV-2 cells infected with GFP-expressing Mm mutants at 48 hpi. The infected cells were monitored by high-throughput, high-content time lapse microscopy for 48 hours. Scale bar, 20 µm.

